# Disrupting IFIT1-STAT2 viral evasion synergy yields live-attenuated flavivirus with enhanced immune priming

**DOI:** 10.64898/2026.06.04.729841

**Authors:** Amporn Suphatrakul, Warangkana Chantima, Pratsaneeyaporn Posiri, Nittaya Srisuk, Rapirat Nantachokechawapan, Nattawadee Panyain, Suppachoke Onnome, Tiwa Rotchanapreeda, Benyapa Thanatummatis, Nopporn Sittisombut, Poonsook Keelapang, Gavin Screaton, Juthathip Mongkolsapaya, Wanwisa Dejnirattisai, Bunpote Siridechadilok

## Abstract

Diverse human viruses deploy a common immune-evasion strategy: viral 2’-O-methylation coupled with STAT2 antagonism. Yet the importance of this coupling and whether targeting the coupled functions offers benefits in vaccine development, remains unknown. Using deep mutational scanning of the dengue NS5 protein, we discovered that the two immune-evasion functions work synergistically to cripple interferon signaling. Disrupting this synergy exposes a reciprocal regulatory relationship between the host proteins IFIT1 and STAT2 that controls both viral restriction and immune cell activation. Dengue viruses with mutations that disrupt the synergy are potently attenuated yet trigger antigen-presenting cell activation that exceeds a licensed dengue vaccine benchmark. Our results establish a generalizable principle: targeting synergies between viral immune evasion mechanisms can simultaneously achieve maximal viral attenuation and innate immune priming. This work provides a rational design framework for developing high-performance vaccines against many human viruses that exploit the IFIT1-STAT2 axis.

**Teaser:** Breaking synergistic viral immune evasion unlocks potent innate immunity for next-generation live-attenuated vaccines.

## INTRODUCTION

The type I interferon (IFN-α and IFN-β) response represents the host’s critical first line of defense against viral infection, serving as the essential bridge between innate sensing and adaptive immunity. To successfully establish infection and evade clearance by immune system, viruses utilize multi-layered strategies to disrupt multiple points along this signaling pathway (*1*). Notably, an evolutionarily conserved counter-strategy across diverse human viruses involves pairing viral 2’-O-methylation (which masks viral RNA from immune sensors and IFIT1; 2’-O-Me) with targeted STAT2 antagonism (which suppresses downstream signaling) (Fig. S1) (*2–33*). Despite the prevalence of this dual-evasion tactic, the mechanistic basis of their co-existence—specifically, whether they operate redundantly or synergistically—and their viability as targets for rational vaccine design have not been established.

Dengue virus (DENV) exemplifies this strategy through its non-structural protein 5 (NS5), a central immune-evasion hub. The NS5 methyltransferase (MTase) domain catalyzes 2’-O-methylation for genomic cloaking, while its RNA-dependent RNA polymerase (RdRp) domain targets STAT2 for proteasomal degradation (*34–36*). Current dengue vaccines, though clinically available, face significant limitations including incomplete cross-serotype protection and variable efficacy in seronegative populations (*37*, *38*). A key challenge is that their attenuation mechanisms are “blind”—they do not account for how natural viruses suppress host immunity, and thus inadvertently undermine the innate immune activation needed to prime robust, durable adaptive responses.

Addressing the mechanistic basis of this functional co-existence requires characterizing the viruses in which the two immune functions are disrupted by mutations across varying infection conditions. However, as overlapping essential functions often densely pack into a single virus gene, this approach faces the challenge of identifying mutations that specifically ablate immune evasion without compromising basal viral replication. Here, we used deep mutational scanning (DMS) across a dengue NS5 gene to systematically identify mutations that specifically compromise viral resistance to type-I interferon. This approach uncovered a critical functional synergy between 2’-O-methylation and STAT2 antagonism and a reciprocal signaling relationship between the host proteins IFIT1 and STAT2. Benchmarking immune analysis of the mutant DENVs showed that the synergistic attenuation of the two functions could afford significantly stronger antigen-presenting cell activation despite being as attenuated as dengue vaccine references. As diverse human viruses possess the coupled immune-evasion functions, our results highlight the importance of the IFIT1-STAT2 signaling axis in immune activation against human virus infections and suggest new target for vaccine development.

## RESULTS

### Deep mutational scanning maps the genetic landscape of type-I IFN resistance by DENV NS5

To identify mutations that specifically affect virus ability to resist type-I IFN, we performed DMS on A549 cells infected with dengue virus library under two conditions, with and without 110 units/ml of type-I IFN (IFN-α2) at the multiplicity of infection (MOI) = 0.1 (Fig. 1A). We used dengue virus serotype 2, strain 16681 (DENV2-16681) library with saturation mutagenesis across all the amino-acid residues of its NS5 gene (*39*). Type-I IFN treatment started 17 hours before infection. Mutant frequencies were assessed two days post infection (dpi). Type-I IFN sensitivity of a variant, a proxy for viral sensitivity to inhibition by type-I IFN, was calculated as the ratio between its fitness score under IFN-α2 = 0 unit/ml and the score under IFN-α2 = 110 units/ml. Pearson correlation coefficients of 0.969 (IFN-α2 = 0 unit/ml) and 0.94 (IFN-α2 = 110 units/ml) were achieved between two replicates of the DMS (Fig. S2A). To select for IFN-sensitizing mutations that minimally affect basal virus replicative ability, IFN sensitivity above 2.5 and DMS fitness scores in Vero cells higher than -3 (which would reduce infectious titer 2 days post infection in Vero cell no less than 1 Log10 relative to the wild-type DENV2 (*39*) (DENV2-WT, blue dots in Fig. S2B)) were used as selection thresholds. These single amino-acid DENV2 mutants did not significantly compromise basal replicative fitness of the DENV2 as evidenced by their titers 2 days post infection (dpi) that were in the range of the titers of DENV2-WT and DENV2-E217A (Fig. S2C). The IFN sensitivity of the selected DENV2 mutants from individual validations is generally consistent with their DMS IFN sensitivity scores (with Pearson correlation of 0.79, Fig. S2D). Combining mutations could increase IFN sensitivity but often at the expense of lowering their basal replicative fitness (titers at 2 dpi in Vero cells) than the corresponding single mutants (Fig. S2C).

**Fig. 1:**
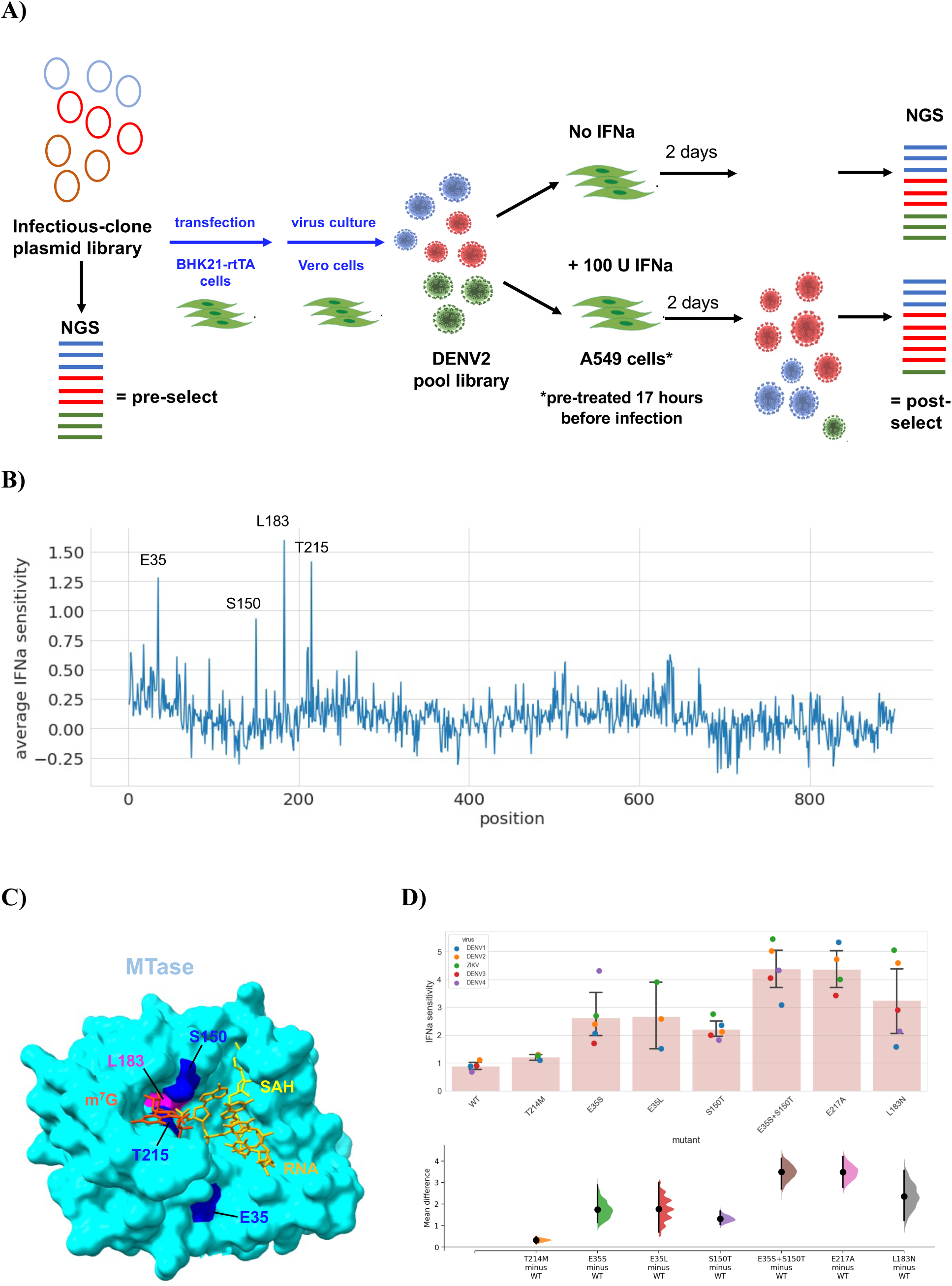
Mapping the genetic determinants of type-I IFN resistance by deep mutational scanning (DMS) of dengue NS5. A) Experimental setup of one replicate of the DMS. B) The DMS-derived average type-I IFN sensitivity per residue across the dengue NS5. C) Structural mapping of the top four type-I IFN sensitive residues E35, S150, L183, and T125 (in blue) on to the ternary complex structure of the MTase domain of DENV3 NS5 (cyan), cap-0 RNA (red), and S-adenosyl-L-homocysteine (SAH) (*32*). D) Type-I IFN sensitivity of WT, T215M, E35S, E35L, S150T, E3S+S150T, E217A, and L183N across DENV1-4 and ZIKV backgrounds (top bar plot). The mean difference for 7 comparisons against the shared control WT are shown in the Cumming estimation plot (*77*). The raw data is plotted on the upper bar plot. On the lower axes, mean differences are plotted as bootstrap sampling distributions. Each mean difference is depicted as a dot. Each 95% confidence interval is indicated by the ends of the vertical error bars. The IFN sensitivity was calculated from the data in Fig. S4B at 500 U/ml for DENV1-4 and at 1000U/ml for ZIKV. With the exception of T215M, the *P* values of the two-sided permutation t-test for all the mutants against WT are less than 0.01.

Strikingly, most IFN-sensitive mutations were concentrated on four residues of the MTase domain: E35, S150, L183, and T215 (Fig. 1B and Fig. S3). S150, L183, and T215 could be mapped onto the active site of the MTase domain and are surrounding the m^7^G cap (Fig. 1C) (*32*). E35 is located toward the edge of MTase active site (Fig. 1C) and is thought to be involved in the interaction with the viral RNA. Alanine mutations of E35, S150 and L183, but not T215, have been shown to reduce cap 2’-O-Me by 40-90% of the wild-type level for WNV MTase (*40*, *41*), suggesting that the identified mutations could affect 2’-O-Me. As E35, S150, L183, and T215 are conserved across flavivirus (*39*), we tested the IFN sensitivity of their mutations in different DENV serotypes and ZIKV backgrounds. Majority of mutations did not reduce the focus sizes of DENV1-4 and ZIKV in Vero cells beyond their standard E217A mutants (except for DENV1-L182N, Fig. S4A), indicating that they did not significantly affect basal virus replication. S150T, E35S, E35S+S150T, and E217A affected viral IFN sensitivity in other virus backgrounds similar to the levels seen in DENV2 (Fig. 1D and Fig. S4B). However, the levels of IFN sensitivity by E35L and L183N were varied across different backgrounds (Fig. 1C). Three levels of IFN sensitivity with DENV2-WT, - S150T/E35S, and -E217A/E35S+S150T were observed (Fig. 1D). Together, these data suggest that E35S, S150T, and E35S+S150T specifically compromise the catalytic function of NS5 MTase in 2’-O-methylation and that they could specifically modulate type-I IFN sensitivity of flavivirus.

### Identification of mutations that specifically ablate STAT2 targeting

Strikingly, no mutations in the residues of the STAT2-binding sites significantly sensitized DENV2-16681 in our DMS condition (Fig. S3). Efforts to test Zika virus (ZIKV) NS5 mutations (G338E+D734A, D734A, and R327A) known to affect STAT2 binding in DENV2 yielded mutant viruses with significantly compromised basal replicative ability (Fig. S5A). To identify STAT2-targeting mutations for DENV2 NS5, we used ZIKV and DENV NS5-STAT2 structures (*35*, *36*) and the NS5 mutant fitness data from DMS in Vero cells (*39*) as the guides to select the mutations for experimental testing. Contact analysis (center-to-center cut-off distance = 5Å) of the four NS5-STAT2 structures showed that the interactions between NS5 and STAT2 were varied (Fig. 2A, top and middle heatmap). NS5 residues such as 334-336, 845-846, 848-851, and 853 are making contacts with STAT2 CCD in all the structures, suggesting that they might be important for the stability of the NS5-STAT2 complexes. However, with the exceptions of residues 334-336, DENV2 do not tolerate mutations at most of these residues well (Fig. 2A, bottom heatmap). Therefore, we selected a subset of mutations that changed the chemical property of the residues with relatively high mutation tolerance and distributed across different STAT2-interacting regions of NS5 such as G21D, K22W, Q26P, P336G, N730D, Q845A, and L853C for the initial round of testing. We also included Q26A, V335I, and V335F as mutations that did not change the chemical properties of side chains significantly. Based on previous study (*36*), D732A+L853A was used as a negative control.

**Fig. 2:**
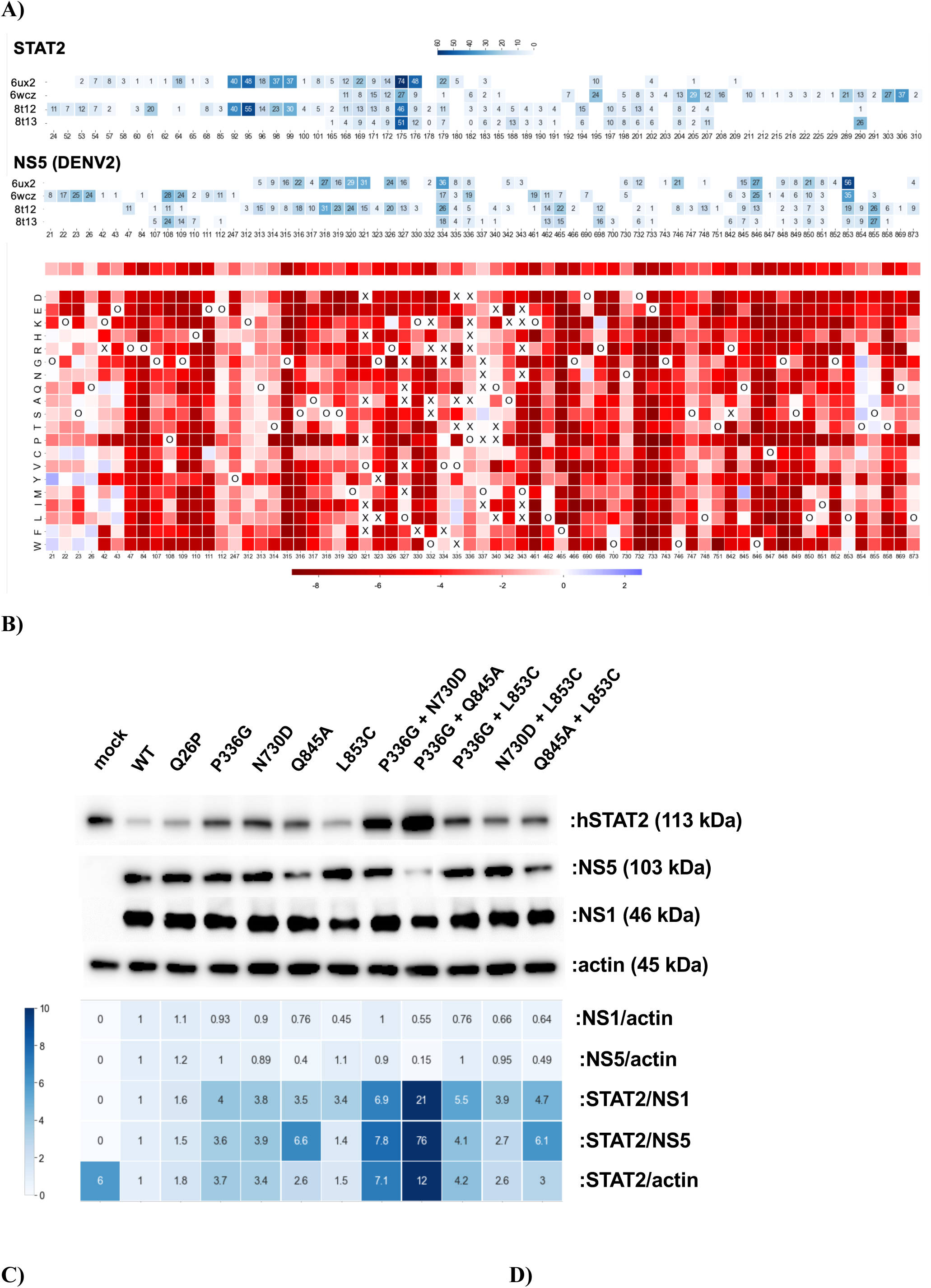
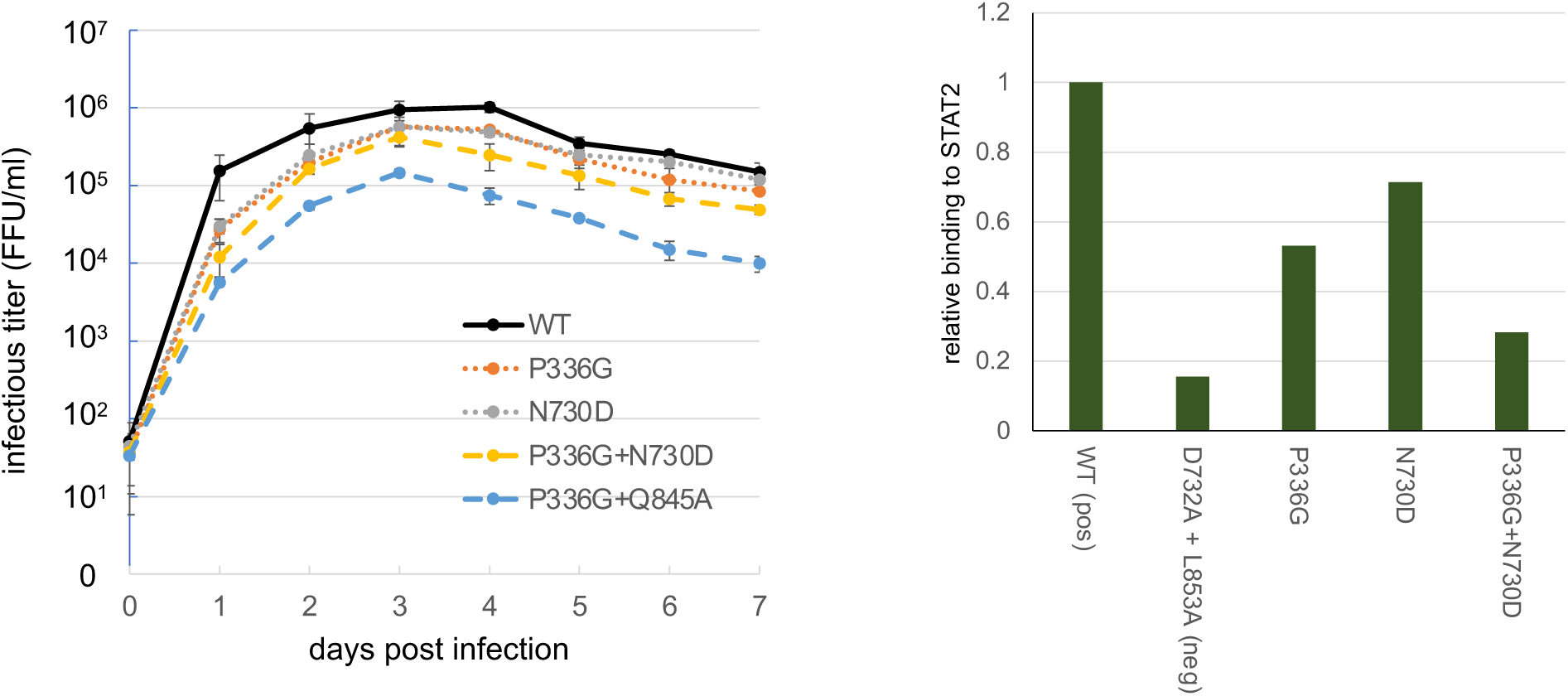
Identification of DENV2 NS5 mutations that specifically compromise its STAT2-targeting function. A) Heatmap of contact analysis of NS5-STAT2 interacting residues from four available NS5-STAT2 complex structures (top and middle, each number represents the numbers of contacts at each residue) with the corresponding mutation fitness data derived from DMS of DENV2 in Vero cells (bottom) (*39*). B) Western-blot analysis of STAT2 degradation during infection by DENV2 with selected NS5 mutations at the STAT2-contacting residues. C) Multi-step replication kinetics of the DENV2-WT, -P336G, -N730D, -P336G+N730D, and - P336G+Q845A in Vero cells infected at MOI = 0.1. D) Relative binding of the NS5 mutants with human STAT2 by co-immunoprecipitation assay.

We used a bimolecular fluorescent complementation (BiFC) assay (*42*) to screen for mutations that disrupt STAT2-NS5 interaction (Fig. S5B). We identified Q26P, P336G, N730D, Q845A, and L853C with similar signal to the D732A+L853A (Fig. S5C). Thus, we constructed DENV2 strains with these mutations (both single and double mutations). All the mutant viruses displayed focus sizes comparable to those of the wild-type DENV2, except for P336G+Q845A that appears significantly smaller (Fig. S5D), confirming that most mutations did not significantly affect basal viral replication.

Next, we tested the ability of the mutant NS5 to degrade STAT2 by assaying for STAT2 level after 2 dpi by Western blot analysis of A549 cells infected with each mutant DENV2 at MOI = 10. To account for potential variations in virus replication, STAT2 levels were normalized by DENV2 NS5 and NS1 level. DENV2 strains with P336G, N730D, and Q845A showed highest STAT2 level in infected A549 cells among the single mutants, 3-5 folds more than the STAT2 level of DENV2-WT infection (Fig. 2C). DENV2 strains with P336G+N730D and P336G+Q845A mutations increased the level of STAT2 in infected A549 beyond the STAT2 level observed in mock-infected cells (STAT2/actin row, Fig. 2C), suggesting improved type-I IFN activation. However, P336G+Q845A showed a decreased ratio of NS1/actin relative to the WT and the P336G+N730D DENV2, suggesting a lower level of virus replication than the two viruses (Fig. 2C). Consistently, the multi-step replication kinetics of the double mutants in Vero cells showed that P336G+Q845A compromised basal replication ability of DENV2 beyond 1 log10, while P336G+N730D did not (Fig. 2D). Thus, high level of STAT2 by P336G+Q845A could be caused by the relatively low level of virus replication that may stimulate STAT2 expression, not the loss of STAT2-targeting activity per se. Quantitative co-immunoprecipitation analysis showed that P336G+N730D almost obliterated NS5 binding to STAT2 while P336G and N730D compromised STAT2 binding to a lesser degree (Fig. 2E), consistent with the STAT2 degradation levels by the DENV2 mutants observed in Fig. 2C. Together, these results showed that P336G+N730D could specifically disrupt STAT2 targeting by DENV2 NS5. An attempt to test the equivalent mutations in other DENV serotypes and ZIKV showed that the mutations did not significantly affect viral basal replication but did not achieve high STAT2 level in infected cells as DENV2-P336G+N730D, suggesting that additional residues determine STAT2-NS5 interaction as STAT2-binding residues among DENVs and ZIKV NS5 are not entirely conserved (Fig. S6).

### Synergy between MTase and the STAT2-targeting functions of dengue NS5

Previous studies have shown that productive replication of DENV2-E217A lacking 2’-O-methylation at the 5’ cap is delayed by around the same time that STAT2 begins to degrade, suggesting the possibility of synergy between viral 2’-O-methylation and STAT2 targeting (*43*, *44*). To test this hypothesis, we constructed a collection of mutant DENV2 that combined mutations from both MTase domain (Fig. 1) and STAT2 binding sites (Fig. 2). While combining E217A greatly compromised viral basal replication (failed to produce virst stock above 10^3^ FFU/ml), MTase mutations identified by our DMS such as S150T could be combined without greatly affecting basal virus replication (Fig. S7A-B), providing suitable strains to test functional synergy. We found that DENV2 strains with only STAT2-targeting mutations (P336G, N730D, and P336G+N730D) had the same IFN sensitivity as the DENV2-WT (Fig. 3A and Fig. S7C), indicating that STAT2 targeting alone could not help virus in breaking through the protection provided by type-I IFN. Adding P336G and N730D failed to enhance IFN sensitivity of the MTase mutants but P336G+N730D could (Fig. 3A and Fig. S7C), indicating that more than four-fold reduction of STAT2 degradation or 50% reduction of STAT2 binding relative to the wild-type virus level would be needed to enhance IFN sensitivity of the DENV2 MTase mutants. Together, these results show that there is a synergy between the viral 2’-O-methylation and STAT2-targeting activities in resisting virus suppression by type-I IFN.

**Fig. 3:**
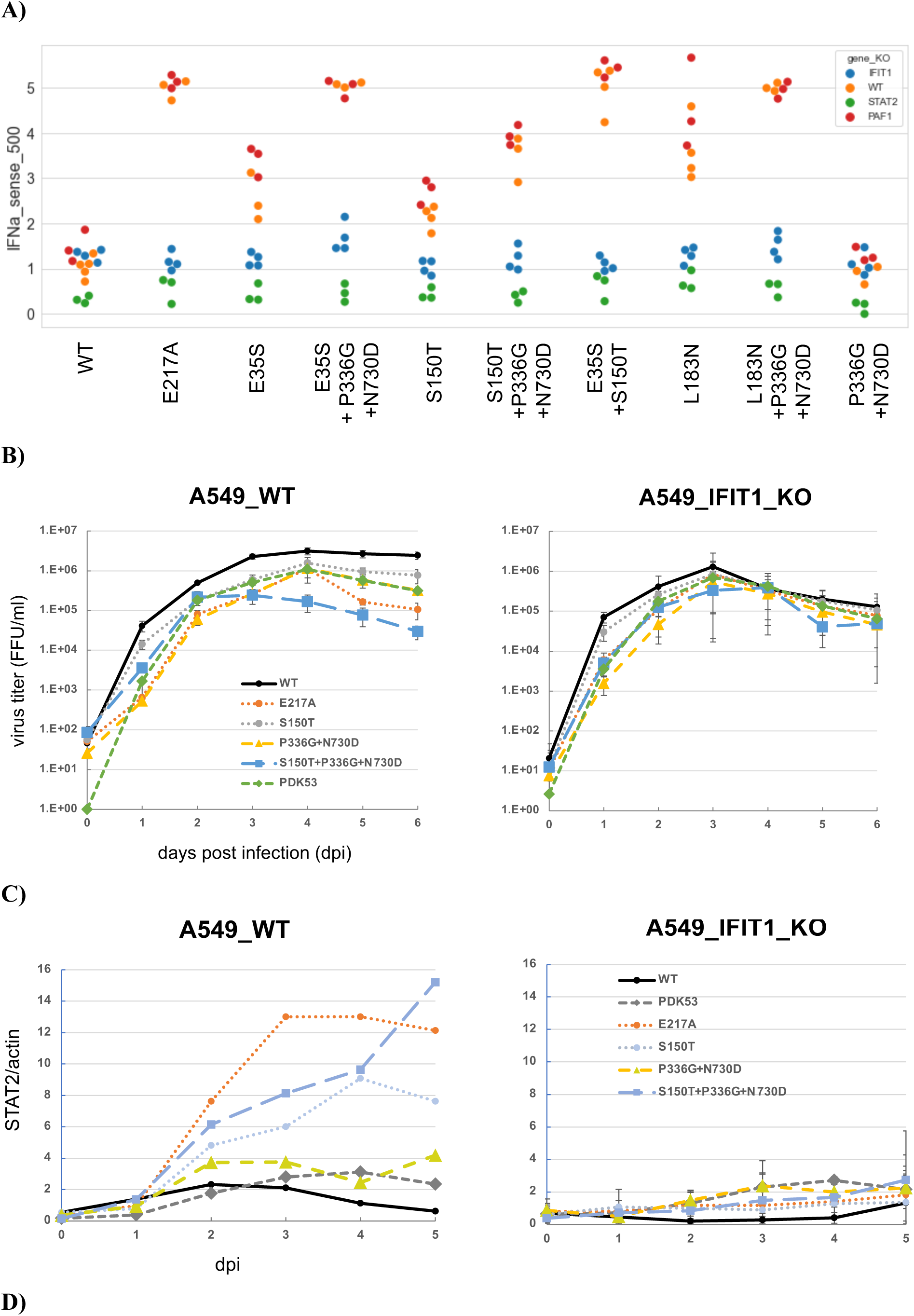
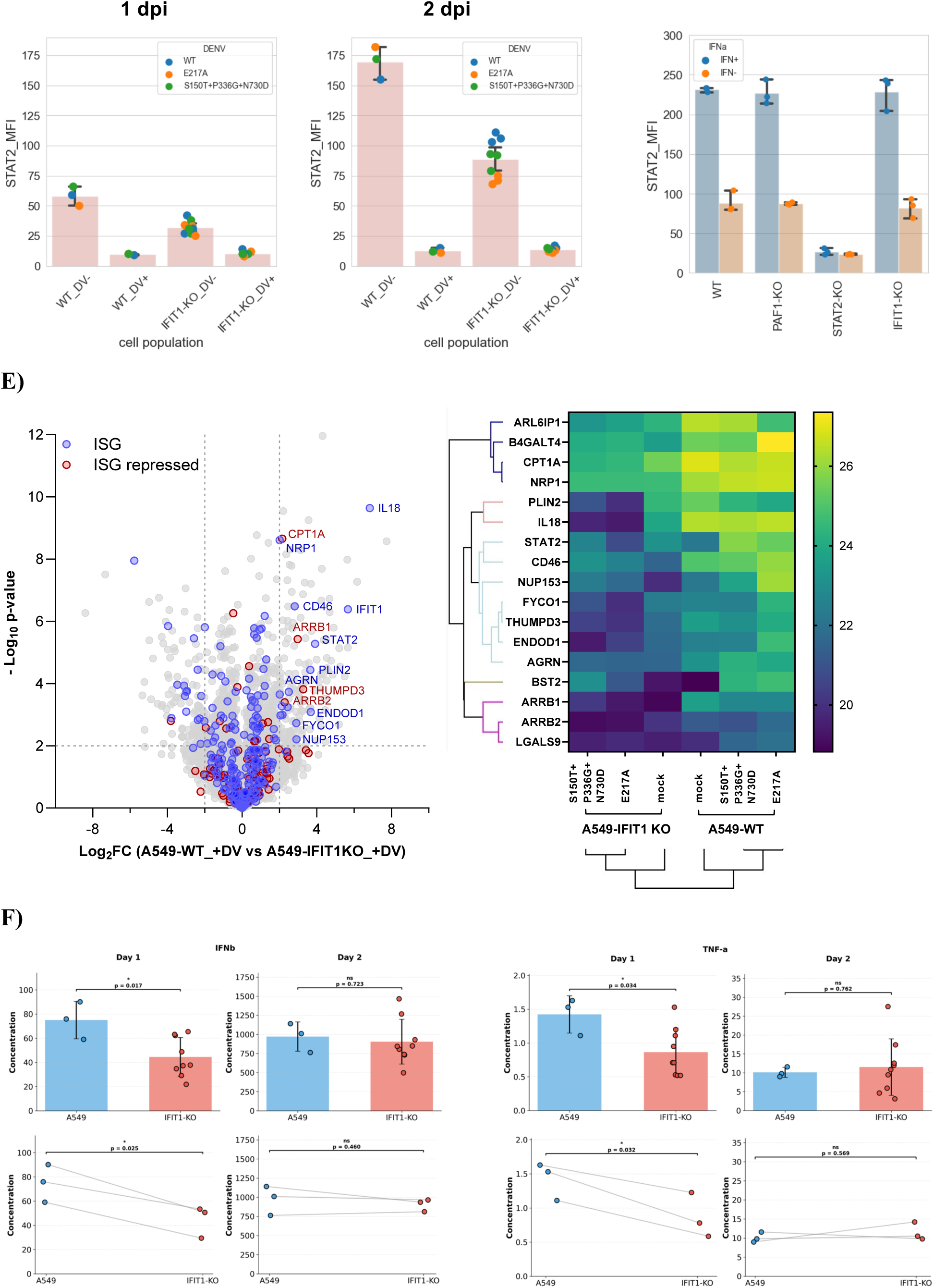
The synergy between viral 2’-O-methylation and STAT2-targeting functions and reciprocal IFIT1-STAT2 signaling relationship. A) Type-I IFN sensitivity of DENV2 with MTase, STAT2-targeting, and combined mutations (MTase+STAT2) in A549-WT and A549-IFIT1-KO, STAT2-KO, and PAF1-KO. The A549 cells were treated with IFN-α2 at 500U/ml for 4 hours before virus infection at MOI = 0.1. The virus titers (FFU/ml) for calculating IFN sensitivity were measured at 2 dpi. B) Multi-step replication kinetics of the DENV2 panel in WT A549 (left plot) and IFIT1-KO A549 (right plot). The A549 cells were infected with DENV2 at MOI = 0.1. C) Western-blot analysis of STAT2 expression level from the multi-step replication kinetics experiment in B). D) Flow cytometry measurement of STAT2 levels (as mean fluorescence intensity, MFI) in the infected and the bystander WT- and -IFIT1-KO A549 at 1 and 2 dpi (left and middle plot) and in the WT- and KO cells treated with type-I IFN at 500U/ml (right plot). E) Proteomic analysis showing differential human protein levels between WT- and IFIT1KO A549 at 2 dpi infected with either DENV2-E217A or DENV2-S150T+P336G+N730D as in B) (left volcano plot). The right heatmap shows the expression profile of ISGs with similar profile to STAT2 in different A549 cells (x-axis). F) Measurement of secreted IFN-β and TNF-α (pg/ml) of A549-WT and -IFIT1-KO cells infected with one of the three viruses, DENV2-WT, - E217A, or S150T+P336G+N730D (MOI = 1) at 1 and 2 dpi. The data points in the top bar plot from each virus were aggregated by cell types. The bottom paired line plots were matched by virus, with the data points in the IFIT1-KO being an average of the measurements from three IFIT1-KO clones. Statistical significance is indicated by brackets above the data sets. Exact p value is provided for each comparison (* ; ns, not significant). Statistical analysis was performed using an unpaired Student’s t-test for the top bar plots and a paired t-test for the bottom line plots.

IFIT1 inhibits flaviviruses by recognizing viral RNA lacking 2’-O-methylation. Using CRISPR-Cas9 knockouts (KO) in A549 cells, we found that IFIT1 KO restored type-I IFN sensitivity of MTase and combined mutants to the wild-type level (Fig. 3A, blue vs. yellow dots), proving IFIT1’s role in the observed synergy. STAT2 KO abolished all IFN sensitivity (Fig. 3A, green vs. yellow dots) while PAF1 KO had no effect (Fig. 3A, red vs yellow dots). Thus, IFIT1 and STAT2 mediate the synergy between viral 2’-O-methylation and STAT2-targeting activities in resisting type-I IFN.

To investigate how viral 2’-O-methylation and STAT2-targeting synergistically support viral replication in immune-competent cells, we compared the multi-step kinetics of DENV2 mutants (S150T, P336G+N730D, and the combined mutant) in wild-type A549 cells. While the viruses showed comparable basal replication in Vero cells (Fig. S7B), the combined mutant (S150T+P336G+N730D) suffered a replication collapse after 2–3 dpi in A549 cells (Fig. 3B). In contrast, the viruses with only one function compromised (S150T and P336G+N730D) continued to replicate efficiently, suggesting these two pathways can partially compensate for each other (Fig. 3B). Notably, the combined mutant and the E217A mutant were more attenuated than the PDK53 benchmark (for details on the PDK53 reference, please refer to Material and Methods). This attenuation was symmetrically reversed in both IFIT1-KO (Fig. 3B) and STAT2-KO cells—but not PAF1-KO (Fig. S8A)—confirming that the viral immune-modulation synergy required to maintain DENV replication in immune-competent hosts also targets IFIT1 and STAT2.

### Identification of reciprocal IFIT1-STAT2 signaling axis

While STAT2 is a known activator of IFIT1, the reverse influence of IFIT1 on STAT2 to account for the observed synergy (Fig. 3A-B) is not known. Western-blot analysis of the A549 cells during the multi-step kinetic analysis in Fig. 3B revealed that DENV2 with MTase mutations (E217A, S150T, S150T+P336G+N730D) induced 3–6 fold higher STAT2 levels than wild-type DENV2, a difference that disappeared in IFIT1-KO cells (Fig. 3C). Analysis by flow cytometry showed that bystander cells expressed higher STAT2 levels than infected cells regardless of virus genetic background (Figure 3D). Flow cytomery and Western blot analysis using anti-NS3 antibodies also confirmed that DENV2-E217A and -S150T+P336G+N730D were stalled in WT cells but spreaded efficiently in IFIT1-KO cells, identifying IFIT1 as a primary barrier to spreading (Supplementary Figures 8B-C). Because IFIT1-KO bystander cells showed lower STAT2 levels than the WT bystanders (Fig. 3D, left and middle bar plots, two-sided permutation t-test, p-values < 0.01) — despite identical STAT2 level when stimulated with the same dosage of IFN-α2 (Fig. 3D, right-handed bar plots, two-sided permutation t-test, p-value = 0.897) —the data suggest that IFIT1 in infected cells enhances the signals sent to neighbouring bystander cells to induce STAT2.

To identify other ISGs influenced by IFIT1, we performed proteomic analysis on WT and IFIT1-KO cells infected with DENV2 with E217A or S150T+P336G+N730D (Fig. 3B). While many classical ISGs (e.g., Mx1, RIG-I) were induced in both backgrounds (Fig. S8D), a specific subset—including IL-18, PLIN2, CD46, and STAT2—showed significantly decreased expression in the absence of IFIT1 (Fig. 3E). These results suggest that IFIT1 does not regulate the entire interferon response, but rather modulates a specific subset of the ISGs during DENV infection.

To pinpoint the specific signaling molecules involved, we performed cytokine profiling of infected cell supernatants by a bead-based multiplex immunoassay. Levels of TNF-α and IFN-β differed significantly between WT and IFIT1-KO cells at 1 dpi (Fig. 3F), while other cytokines remained unchanged (Fig. S8E-8F). Given that STAT2 induction is directly proportional to type-I IFN dosage (Fig. S7G), these results suggest that IFIT1 could regulate the level of STAT2 in bystander cells through a combination of cytokines (consisting of at least TNF-α and IFN-β) from infected cells. Collectively, these results establish an intercellular reciprocal signaling axis where IFIT1 governs the paracrine induction of STAT2 (and a subset of ISGs) in bystander cells to limit viral spread.

### Attenuation of the synergistic immune-evasion functions enhances innate immune response in antigen-presenting cells

To understand how viral 2’-O-methylation and STAT2-targeting functions affect professional antigen-presenting cells (APCs) that are target cells of DENV (*45–48*), we analyzed 14 cellular/infection markers and 12 key cytokines/chemokines across human monocyte-derive dendritic cells (MDDCs), M1, and M2 macrophages using 10 DENV2 mutants (243 measurements). Principal Component Analysis (PCA) and K-means clustering showed that while WT, PDK53 reference, MTase, and STAT2-targeting mutants formed distinct, predictable groups, the combined mutants—particularly S150T+P336G+N730D—exhibited unique immune profiles (Fig. 4A). This separation suggests that the synergy between 2’-O-Me and STAT2-targeting functions fundamentally alters how APCs perceive and respond to the virus.

**Fig. 4:**
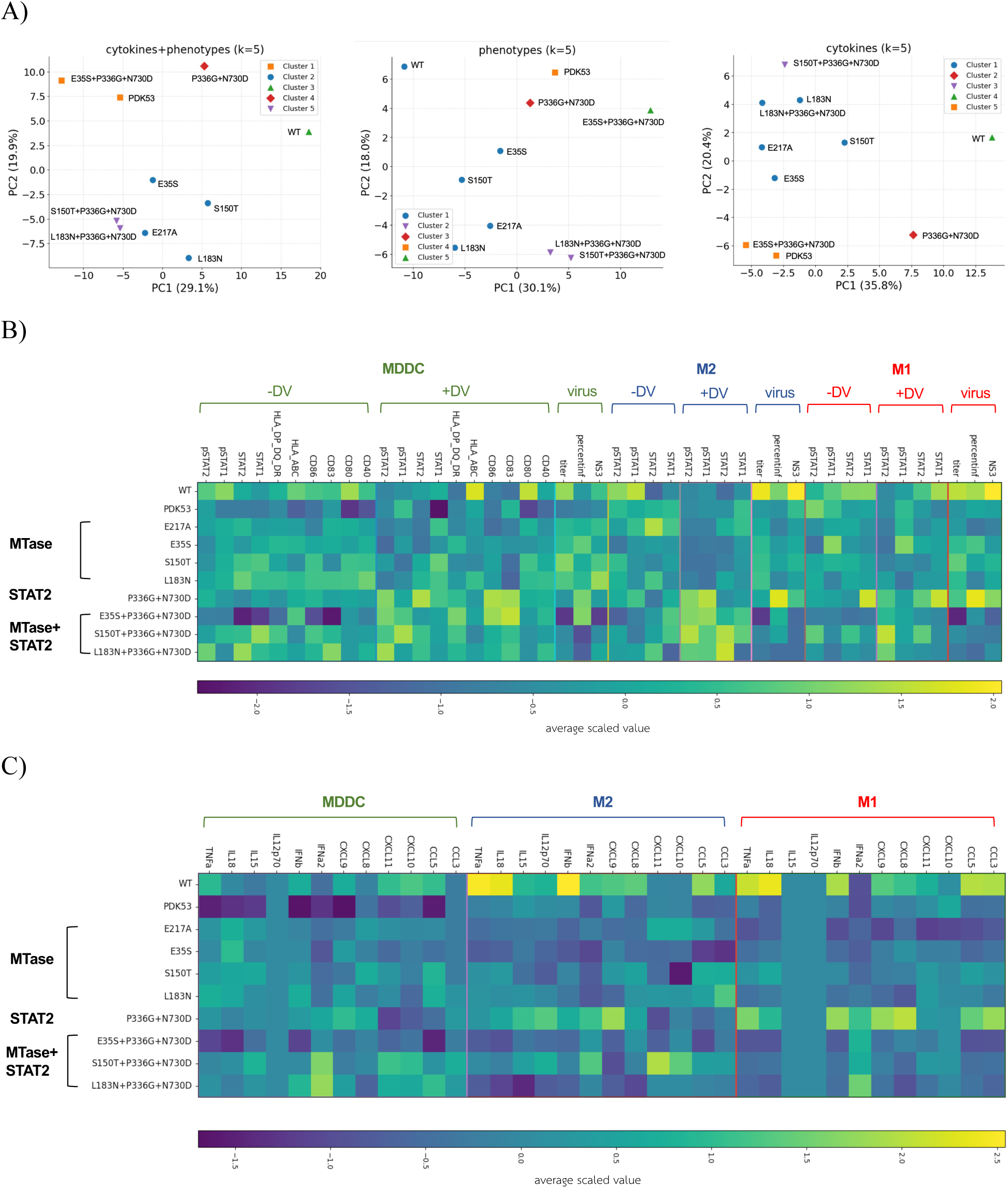
Synergistic attenuation mutations targeting dengue NS5 2’-O-methylation and STAT2-targeting functions could enhance the activation of human antigen-presenting cells. A) Principal component analysis and K-mean clustering of immune profiles of the DENV2 panel. B) Heatmap of phenotype (+/-DV) and infection (virus) measurements in MDDC, M1, and M2 across the DENV2 panel. DV+ = infected cells, DV- = bystander cells, virus = infection parameters. C) Heatmap of cytokine and chemokine measurements of MDDC, M1, and M2 infected by the DENV2 panel.

Compromising the STAT2-targeting function of NS5 (P336G+N730D) significantly increased the activation of infected MDDCs and M2 macrophages, characterized by elevated pSTAT2 and MDDC maturation markers CD83 and CD86 (Fig. 4B and Fig. S9A-D). Interestingly, M1 macrophages required the full synergistic mutations (S150T/L183N+P336G+N730D) to reach a peak activation level (Fig. 4B and Fig. S9C). In contrast, the PDK53 reference and single MTase mutants failed to enhance these markers, implicating NS5-mediated STAT2 inhibition as the primary mechanism DENV used to dampen APC activation (Fig. 4B). Cytokine profiling further highlighted the impact of synergistic mutations, as the S150T/L183N+P336G+N730D mutants uniquely enhanced IFN-α2 production in MDDCs (Fig. 4C and Fig. S9E). While the PDK53 reference and MTase mutants generally reduced pro-inflammatory cytokines (TNF-α, IL-18) in macrophages, the combined mutants maintained the cytokines at higher levels (Fig. S9F-G). Notably, S150T+P336G+N730D specifically boosted T-cell attracting chemokine CXCL11 in M2 macrophages, a profile not seen with other DENV2 strains (Fig. S9F-G). In summary, perturbing the two viral IFN-antagonistic functions enhances APCs activation and cytokine induction in a cell-specific manner. Activation of M1 appeared least sensitive to the perturbations while MDDC and M2 were similarly sensitive, highlighting the distinct immune response across the APCs during DENV infection.

Despite its major role in dampening APC activation (Fig. 4B), disrupting STAT2-targeting function alone was not sufficient to attenuate virus replication in APCs to the same level as observed in the PDK53 reference. DENV2 with P336G+N730D mutation replicated at the levels similar to the DENV2-WT (Fig. 4B and Fig. S9H-J). In contrast, DENV2-S150T/L183N+P336G+N730D replicated at similar or lower levels but activated infected APCs better than the PDK53 reference and E217A (Fig. 4B-C and Fig. S9H-J). Together, these results identify DENV2-S150T+P336G+N730D as high-potential candidates for vaccine development.

In addition to illuminating the role of STAT2-IFIT1 signaling axis, our results show both common and distinct signaling responses across the APCs during DENV infection. DENV2 induced stronger STAT1/2 expression and their phosphorylation in bystander cells than in infected cells across the three APCs (Fig. S10A-C). While certain chemokines (CXCL10, CXCL11, CCL5) were universally upregulated during DENV2 infection, cytokines like IFN-α2, CXCL8, IL-18, CXCL9, CCL3, IFN-β, TNF-α, and IL-15 showed cell-specific patterns, with higher upregulation in MDDCs than in macrophages (Fig. S10D-G).

## DISCUSSION

By applying DMS to DENV2 under varying immune pressures (A549 IFN-α2 vs. Vero cells), we revealed a potent functional synergy between 2’-O-methylation and STAT2-targeting activities (Fig. 1-3). Crucially, disrupting this synergy challenges the traditional model of IFIT1 function. While IFIT1 is canonically viewed as a downstream effector of the type-I IFN response (*49–51*), our data suggest it acts as an amplifier of TNF-α and type-I IFN secretion rather than a negative regulator (*52*). Importantly, this reinforcement loop operates independently of the IFIT3-MAVS-TBK1 pathway [27–29], as IFIT3 levels remained unaffected during DENV infection in IFIT1-knockout cells (Fig. 3E and Fig. S8D). Thus, our results provide a mechanistic explanation of how IFIT1 can function as a central innate immune bottleneck (*53*) that orchestrates a broader antiviral state during infection through reciprocal IFIT1-STAT2 signaling axis (Fig. 3).

By targeting this synergistic bottleneck, our results highlight a pivotal trade-off in live-attenuated vaccine (LAV) design: the necessity to maximize innate immune activation without crippling the basal replicative fitness required for antigen production. Even though both DENV2-S150T+P336G+N730D and E35S+P336G+N730D were attenuated by targeting the synergistic functions, the one with higher basal replicative fitness (S150T+P336G+N730D) generated a superior immune-cell activation profile compared to its weaker counterpart (E35S+P336G+N730D), the E217A mutant, and the PDK53 benchmark (Fig. 4 and Fig. S9). Because robust activation and maturation of primary human antigen-presenting cells (APCs) provide the essential co-stimulatory bridge to durable adaptive immunity, this strategy addresses the critical shortcoming of historical, “blind” attenuation. Aligning with recent studies in Zika (*54*, *55*) and influenza (*56*), our data establish that precisely tuning immune-antagonistic functions—rather than simply weakening basal replicative ability of a virus as used in conventional LAV development—is the optimal route for eliciting potent immune priming.

The synergy between 2’-O-methylation and STAT2 antagonism likely evolved to navigate the specific host-restriction landscapes of the human immune system. Given that both IFIT1 and STAT2 evolved rapidly across mammalian species (*57*, *58*), this dual-evasion strategy could confer a distinctly species-specific advantage for human infection. This human-centric adaptation underscores the critical value of using primary human APCs to model innate immunogenicity, as standard small animal models often fail to accurately recapitulate flavivirus STAT2 antagonism (*59*). Because diverse human-tropic viruses utilize this coupled evasion strategy (Fig. S1), disrupting this synergistic node could provide a rigorous strategy for rationally attenuating a wide range of emerging human pathogens.

Finally, viral proteins are often highly pleiotropic, featuring overlapping structural and immune-modulating domains that confound traditional mutagenesis. We demonstrate that integrating DMS, structural modeling, and host genetic perturbation can systematically untangle these complex functional interactions. Looking forward, this high-resolution mapping approach offers a direct solution to the enduring challenge of multivalent dengue vaccine formulation. By enabling developers to precisely dial the replicative fitness and intrinsic immunogenicity of individual strains, this paradigm could resolve the unbalanced baseline immunity that has historically plagued tetravalent dengue vaccines.

## MATERIALS AND METHODS

### Ethical statement and biosafety standard

This study was reviewed and approved by the Institutional Review Board (IRB) of the Faculty of Medicine, Siriraj Hospital, Mahidol University (COA no. Si 221/2024). Written informed consent was obtained from all individual participants prior to their inclusion in the study. All experimental protocols involving recombinant DNA and flavivirus (DENV and ZIKV) were reviewed and approved by the Institutional Biosafety Committee (IBC) of the Faculty of Medicine, Siriraj Hospital, Mahidol University (SI2024-007, SI2023-069, SI2018-007). All procedures involving infectious viruses were performed in a certified Biosafety Level 2 (BSL-2) facility.

### Cell culture

Vero cells were maintained in DMEM (low glucose) supplemented with 10% heat-inactivated fetal calf serum (HI-FCS), penicillin (100 units/ml), streptomycin (100 μg/mL), and L-alanyl-L-glutamine (GlutaMAX, 2mM) (DEM10). C6/36 cells were maintained in Leibovitz’s L-15 supplemented with 10% HI-FCS, penicillin (100 units/ml), streptomycin (100 μg/mL), and L-alanyl-L-glutamine (GlutaMAX, 2mM) (L10). A549 cells were maintained in RPMI1640 supplemented with 10% HI-FCS, penicillin (100 units/ml), streptomycin (100 μg/mL), and L-alanyl-L-glutamine (GlutaMAX, 2mM), and 1x MEM non-essential amino acids (R10-AA). Vero and A549 were cultured at 37°C, 5% carbon dioxide, with relative humidity between 90-95%, while C/36 were culture at 28°C. Gesicle producer 293T cells were maintained in DMEM (high glucose) supplemented with 10% heat-inactivated fetal calf serum (HI-FCS), penicillin (100 units/ml), streptomycin (100 μg/mL), and L-alanyl-L-glutamine (GlutaMAX, 2mM) (DEM10-HG).

The preparation of human dendritic cells and macrophages started with the isolation of peripheral blood mononuclear cells (PBMCs) from heparinized whole blood obtained from healthy donors using Ficoll-Hypaque density gradient centrifugation and followed by the isolation of CD14+ monocytes from PBMCs using MACS CD14 MicroBead mAbs (Miltenyi Biotec). The isolated CD14+ cells were cultured in RPMI 1640 supplemented with 10% FCS (R10), 25 ng/ml recombinant human GM-CSF (PeproTech), and 50 ng/ml recombinant human IL-4 (PeproTech) for 6 days in a 37°C incubator with 5% CO2 to differentiate the CD14+ monocytes into immature monocyte-derived dendritic cells. We then assessed the phenotype and purity of the cells by examining the expression of CD14 and CD209 on the cell surface by flow cytometry using the antibodies specific to both markers. Macrophages were then derived from monocytes. Based on nomenclature that utilizes other stimuli for macrophage polarization (i.e., IFN-γ and LPS for M1, or IL-4 and IL-13 for M2), following differentiation with GM-CSF or M-CSF, both cell types would be classified as equivalent to the M0 subtype. To generate M1-like macrophages, CD14+ cells are cultured in R10 containing 50 ng/ml recombinant human GM-CSF (PeproTech), whereas M2-like macrophages are cultured in R10 containing 50 ng/ml recombinant human M-CSF (PeproTech) for 5 days in a 37°C incubator with 5% CO_2_. Both macrophage types could be infected by dengue viruses without requiring antibody-dependent enhancement. We confirmed the subtypes using a panel of specific antibodies (BD): M1 exhibited relatively high expression levels of MHC II (HLA-DR) and co-stimulatory molecules such as CD80 and CD86, while M2 expressed CD163 and CD206 molecules. The primary cells were maintained at 37°C, 5% carbon dioxide, with relative humidity between 90-95%.

### Viruses

All the DENV1-4 and ZIKV wild-type and mutant strains used in this study were generated from infectious-clone plasmids. The plasmids were transfected into BHK21-rtTA3 cells using Lipofectamine 2000, following a previously reported protocol (*60*). Subsequently, the cells were cultured in ISF-1 medium supplemented with 1 µg/ml doxycycline for an additional three days to induce virus production. The viruses were then propagated in Vero cells maintained in ISF-1 medium for approximately 2–3 days prior to harvesting. Cell debris was removed by centrifugation at 4,000 rpm and 4°C for 20 minutes. The clarified supernatant containing viral particles was supplemented with HI-FCS to a final concentration of 20% and stored at -70°C. Virus stocks used in experiments with human dendritic cells and macrophages were propagated for another round in C6/36 cultured at 28°C using L-15 supplemented with 2% FCS and harvested between 2-3 days post infections. The viruses in the culture media were harvested by centrifugation at 10,000 xg, at 4°C for 20 hours. The pellets were resuspended with L10 and stored frozen at - 70°C until use. We used 16681-3pm, a recombinant strain harboring the attenuating point mutations of strain PDK53, as the PDK53 reference (*61*). 16681-3pm displays key attenuation phenotypes similar to the original PDK53 such as the pinpoint plaque sizes, reduced viremia, and immunogenicity in nonhuman primates in the context of chimeric prM-E strains across the four dengue serotypes (*61–65*). The virus stocks were sequence-verified by Sanger sequencing. Infectious viral titers were determined using a foci forming assay utilizing the 4G2 monoclonal antibody.

### Virus titer determination by foci assay

Infectious titers of DENV were quantitated by foci assay with 4G2 monoclonal antibody according to the previous report (*66*). The 96-well plates were imaged using CTL S6-Ultimate-V Analyzer (CTL Analyzers).

### Plasmid construction

The infectious clone plasmids of DENV1-4 and ZIKV with an intron insert will be detailed elsewhere (Suphatrakul et al., manuscript in preparation). Briefly, the viral RNA was extracted from virus stocks using QIAamp viral RNA extraction kit according to the manufacturer’s protocol. cDNA was generated using Superscript III according to the protocol in Siridechadilok et al. 2013(*66*). cDNA was used as the template to generate DNA fragments for assembly by PCR amplification using Phusion DNA polymerase. PCR fragments were assembled with vector by Gibson assembly (*67*). The assembly reaction was cleaned up by AMPure XP magnetic bead (Beckman Coulter) and electroporated into DH5a. For this study, DENV1 strain KDH, DENV2 strain 16681, DENV3 strain v1043, DENV4 strain H241, and ZIKV strain SV0010 were used.

The NS5 gene of DENV2-16681 was sub-cloned from the infectious-clone plasmid, fused with a C-terminal HA tag sequence, and inserted downstream of the inducible CMVtight promoter into a lentiviral plasmid containing an eGFP expression cassette as a marker. Similarly, the human STAT2 gene was sub-cloned from pUNO1-hSTAT2 (InvivoGen), fused with a C-terminal FLAG tag sequence, and cloned into a lentiviral plasmid under the control of the inducible CMVtight promoter, utilizing an mCherry expression cassette as a marker. The HA and FLAG tags were utilized for pull-down assays using anti-HA and anti-FLAG antibodies, respectively. The plasmids were cloned and propagated in NEB-Stbl.

The plasmid for expressing single guide RNA (sgRNA) for CRISPR-Cas9 was constructed from pC0043 plasmid (Addgene #103854) by double cleavage with BsbI and XhoI to replace the PspCas13b-crRNA with that of annealed primers containing Cas9 sgRNA. Bsb1 site was introduced between the U6 promoter and the scaffold for cloning a spacer sequence. A spacer sequence was cloned by ligation of the phosphorylated annealed primers using T4 DNA ligase. The plasmids were cloned and propagated in DH5a.

### Deep mutational scanning

We used the DENV2 infectious-clone plasmid library as described previously (*39*). The infectious-clone plasmid library was transfected into BHK21-rtTA3 treated with 1μg/ml doxycycline during seeding and maintained after transfection. The media was harvested 3 days post transfection and transferred to infect confluent Vero cells. The final DENV2 pool libraries were harvested twice at 3 and 5 days post infection. Each DENV2 pool library was precipitated with 10% polyethylene glycol 8000 (PEG8000) overnight at 4°C and harvested by centrifugation at 3200xg, 4°C, 30 minutes. The pellet was resuspended in 1xPBS and supplemented with 20% FBS for storage. A549 cells were infected with virus library at MOI = 0.1 for two hours. The cells were then washed three times with 1xPBS before being further cultured in RPMI1640 + 10% heat-inactivated fetal calf serum at 37 °C for 48 hours before collecting the released virus from the cells in the cell culture medium for further analysis. Testing in A549 cells was done in two conditions. In the first condition, A549 cells were not exposed to interferon alpha in the culture medium for the entire duration of the experiment. In the second condition, A549 cells were exposed to 110 U/ml of interferon alpha in the culture medium starting at 17 hours before virus infection and continued to receive 110 U/ml of interferon alpha throughout the experimental period. Viruses generated and released in cell culture medium were precipitated with PEG8000 at 10% concentration overnight at 4 °C before centrifugation at 3200 xg, 4 °C, 30 min. The pellet was dissolved with 1x PBS by vortex or aspirated with a pipette. The resuspended pellet was supplemented with the final concentration of 10% heat-inactivated fetal bovine serum for storage at -70°C. Half of the collected virus was then extracted with a viral nucleic acid extraction kit II (Geneaid) and the RNA was converted into complementary DNA (cDNA) using the ImProm-IITM reverse transcription system (Promega). Polymerase chain reaction (PCR) was used to extract the location of the NS5 mutation for sequencing. The PCR product was quantified with Nanodrop and Qubit. From all the 23 batches of virus library pools, equal amount of the PCR product (in ng) from each pool was then mixed into a single tube for deep sequencing with Illumina Novaseq 150bp paired-end reads.

DMS data analysis was performed as described previously (*39*). We used DiMSum wrapper to pre-process the fastq data using default settings (*68*). We generated the final count of each mutant using a custom python script. We set the input read-count thresholds to 60, 25, and 25 for mutants with the Hamming distances of 1, 2, and 3 nucleotides, respectively. From the read counts, we re-created the vsearch.unique files of the input and the outputs with full-length NS5 sequences for fitness score calculation with normalization and error model fitting by STEAM analysis in DiMSum. Mutant read counts in the infectious-clone plasmid library were used as the input while the mutant read counts from the DENV2 virus library generated from A549 were used as the output for calculating fitness scores of the mutants under +/- IFN-α2. The IFN sensitivity of the NS5 mutants were visualized using WebLogo plot created by dms2_logoplot (Fig. S3) (*69*).

### Multi-step replication kinetics

Vero cells were infected with the DENV2 at MOI = 0.1 in DMEM supplemented with 2% HI-FCS (DMEM-2) for 2 hours and washed with 1xPBS twice before re-supplmented with fresh DMEM-2. A549 cells were incubated with DENV2 at MOI = 0.1 in RMPI1650 supplemented with 10% HI-FCS (R10) for 2 hours and washed with 1xPBS twice before re-supplemented with fresh R10. The infected cells were maintained at 37°C, 5% carbon dioxide, with relative humidity between 90-95%. The culture was swirled to mix before harvesting the media at each time point. The media were clarified by centrifugation at 4,000 rpm and 4°C for 20 minutes. The clarified supernatant containing viral particles was supplemented with HI-FCS to a final concentration of 20% and stored at -70°C. The infectious titers were determined using foci assay as described above.

### Measuring interferon sensitivity

Each mutant virus was subjected to the tests in Vero cells and in A549 cells to measure its basal fitness and interferon sensitivity. Vero cells were infected with the virus series at MOI = 0.1 for two hours. The cells were then washed with 1xPBS three times before being cultured in MEM + 2% HI-FCS (MEM-2) at 37 °C for 48 hours before harvest of culture media for further analysis. The media were clarified by centrifugation at 4,000 RPM (or 2576xg, Sorvall ST 8R/Thermo scientific) and 4°C for 20 minutes. The clarified supernatant containing virus particles was supplemented with HI-FCS to a final concentration of 20% and stored at -70°C. A549 cells were infected with the virus at MOI = 0.1 for two hours. The cells were then washed with 1xPBS three times before being cultured in R-10 at 37 °C for two days before harvest of culture media for further analysis. Testing in A549 cells was performed in two conditions. In the first condition, A549 cells were not exposed to IFN-α2 for the entire duration of the experiment. In the second condition, A549 cells were exposed to 100 U/ml IFN-α2 in the culture medium starting either at 17 hours or 4 hours before virus exposure and continued to receive the same dose of IFN-α2 throughout the experimental period. Viruses generated and released in the cell medium were measured using foci assay with 4G2 monoclonal antibody. Interferon sensitivity was calculated as the logarithm of the ratio between the infectious titer of the mutant virus derived from A549 cells that did not receive IFN-α2 to that of the mutant viruses derived from A549 cells that received IFN-α2.

### Analysis of STAT2 degradation by Western-blot analysis

To evaluate the ability of the virus to degrade STAT2, A549 cells seeded in 24-well plates were infected with DENV1-4 at MOI = 10 or ZIKV at MOI = 5 for four hours. The cells were then washed three times with 500 µl of 1x PBS and maintained in R-10 supplemented with 1x MEM non-essential amino acids for an additional 48 hours. For mock infection, cells were treated with R-10 in the absence of the virus. Cells were washed twice with ice-cold 1x PBS and harvested using 2x Laemmli sample buffer for Western blot analysis. DENV2 NS5 protein levels were detected using the mouse monoclonal antibody GT353 (Thermo Fisher). DENV1-4 and ZIKV NS1 levels were detected with the mouse monoclonal antibody 1F11. STAT2 levels were detected using the rabbit monoclonal antibody D9J7L (Cell Signaling). Lysate loading was normalized against beta-actin levels, which were detected using the mouse monoclonal antibody 8H10D10 (Cell Signaling). Membranes incubated with the aforementioned primary antibodies were subsequently probed with HRP-conjugated secondary antibodies (1:4,000 dilution) for 1 hour. Following three washes with 10 ml of TBS-T for 5 minutes each, the blots were visualized using an Amersham ImageQuant 800 Biomolecular Imager. Band intensities were analyzed using ImageQuant™ TL Version 8.2 software (GE).

### Bimolecular fluorescent complementation (BiFC) assay

To assess NS5–STAT2 interactions, we employed a split-YFP system using pBiFC-VN155(I152L) (Addgene #27097) and pBiFC-VC155 (Addgene #22011). Structural modeling of the NS5-STAT2 complex (PDB: 6WCZ) and AlphaFold2 predictions of the disordered STAT2 C-terminus informed our tagging strategy. DENV2 (strain 16681) NS5 and human STAT2 cDNAs were cloned via Gibson assembly to generate N- and C-terminal fusions with VN and VC domains.

293F-rtTA3 cells were seeded at 200,000 cells/well in 24-well plates (DMEM, 10% FBS, 1x NEAA) 24 h prior to transfection. Eight fusion combinations (3 µg total DNA; 1:3 DNA:PEI ratio) were screened to determine the optimal orientation. Based on maximum fluorescence 48 h post-transfection (fluorescence microscopy), the N-terminally tagged NS5 (VN-NS5) and C-terminally tagged STAT2 (STAT2-VC) pair was selected. The system was validated using the binding-deficient NS5 D732A/L853A mutant as a negative control.

Specific NS5 mutations (G21D, K22W, Q26A, Q26P, P336G, N730D, Q845A, L853C, V335A, and V335F) were introduced via site-directed mutagenesis. Cells were transfected and harvested 48 h post-transfection, then fixed and analyzed via flow cytometry (BD Fortessa). Mutations resulting in fluorescence signals comparable to the D732A/L853A control were identified as disruptors of the NS5–STAT2 interaction.

### Co-immunoprecipitation of NS5-STAT2

293F-rTtA3 cells were cultured in DMEM-10 supplemented with 1x MEM non-essential amino acids in 6-well plates. Cells were initially seeded at a density of 8×10^5^ cells in 2 ml volume and incubated at 37°C with 5% CO_2_ for 20–24 hours prior to transfection. Before transfection, half of the culture medium (1 ml) was replaced, and doxycycline was added to a final concentration of 1 µg/ml. The transfection mixture was prepared by combining equal amounts of pLenti-mCh-hSTAT2-FLAG and pLenti-GFP-NS5-Mutant-HA plasmids (1.5 µg per plasmid, concentration determined by Qubit) in serum-free DMEM to a final volume of 100 µl. Simultaneously, polyethylenimine (PEI) was prepared by mixing 9 µl of PEI (1 mg/ml) with 91 µl of serum-free DMEM, resulting in a DNA:PEI ratio of 1:3 (3 µg DNA total to 9 µg PEI). The DNA and PEI solutions were combined (total volume 200 µl), incubated at room temperature for 15 minutes, and added dropwise to the cells. The cells were then returned to the incubator at 37°C with 5% CO2. On the following day, the medium was replaced with 2 ml of fresh medium containing 1 µg/ml doxycycline. At 48 hours post-transfection, cells were washed with PBS, dissociated into single cells using 2 mM EDTA, and neutralized with culture medium. Cells were transferred to 1.5 ml tubes and counted. A total of 1×10^6^ cells were centrifuged at 800×g for 3 minutes at 4°C, washed with 500 µl of PBS, and centrifuged again. The supernatant was removed, and the cell pellet was resuspended in 250 µl of cold NP40-lysis buffer (0.25% NP-40, 50 mM Tris pH 7.4, 150 mM NaCl, 5 mM EDTA, 10% glycerol, and 1X Protease inhibitor cocktail). Lysis was performed by gentle pipetting (30 cycles), followed by rotation for 30 minutes at 4°C. Lysates were clarified by centrifugation at 14,000×g for 10 minutes at 4°C.

For immunoprecipitation, 220 µl of the clarified lysate was transferred to a new 1.5 ml low-binding tube. Anti-FLAG monoclonal antibody (1 µl of 1 µg/µl, Sigma F1804) was added, and the mixture was rotated for 1 hour at 4°C. Subsequently, 10 µl of Protein G Sepharose™ 4 Fast Flow slurry was added, and the mixture was rotated overnight at 4°C. The following day, the resin was pelleted by centrifugation at 1,000×g for 1 minute at 4°C. The supernatant was discarded, and the resin was washed three times with 1 ml of NP40-lysis buffer. After the final wash, all supernatant was removed, and the resin was resuspended in 40 µl of 2X Laemmli buffer. The samples were boiled at 95°C for 5 minutes and centrifuged at 1,000×g for 5 minutes at 25°C. The eluate was transferred to a new tube and stored at -20°C for Western blot analysis.

Samples prepared from the co-immunoprecipitation were loaded onto two 4–12% gradient NuPAGE Bis-Tris gels (Invitrogen). Proteins were transferred to Immobilon PVDF membranes using an ECL Semi-dry Blotter (GE Healthcare) at 80 mA for 1 hour. Membranes were blocked with 5% skim milk in TBS-T for 1 hour. One membrane was probed with anti-HA antibody (HA-Tag (6E2) Mouse mAb #2367, Cell Signaling; 1:1,000 dilution) and the other with anti-FLAG antibody (DYKDDDDK Tag Antibody-Rabbit #2368, Cell Signaling; 1:1,000 dilution) in 3% BSA in TBS-T overnight. Membranes were washed three times with 10 ml TBS-T (5 minutes each) and incubated with HRP-conjugated secondary antibody (1:4,000 dilution) for 1 hour. Following three additional washes with 10 ml TBS-T, signals were detected using an Amersham ImageQuant 800 biomolecular imager, and band intensities were analyzed using ImageQuant™ TL version 8.2 (GE).

### Gene knock out by CRISPR-Cas9

To deliver CRISPR-Cas9 into A549 cells as ribonucleoprotein (RNP) complex, we utilized the Cas9-eVLPs platform (*70*). Cas9-eVLPs targeting IFIT1, STAT2, and PAF1 were produced by transient transfection of Gesicle Producer 293T cells. The 293T cells were seeded at a density of 18x10^6^ cells per 150mm dish. After 20–24 hours, cells were transfected with a mixture of plasmids expressing VSV-G (1,440 ng), MLVgag–pro–pol (12,150 ng), MLVgag–Cas9 (4,050 ng), and two pC0043-sgRNAs (15,840 ng) using PEI. After 40–48 h post-transfection, producer cell culture media was harvested and centrifuged for 10 min at 2000xg to remove cell debris. The clarified supernatant was then filtered through a 0.45-mm PVDF filter. The eVLPs were precipitated and then pelleted using the PEG Virus Precipitation kit (ab102538 ; Abcam). The pellet was resuspended with 200-400 µl 1xPBS and stored at -80°C until use. 50-100 µl Cas9-eVLPs were used to transduce A549 cells in 24-well plate (plated at density of 75,000 cells per well). After 20–24 hour seeding, Cas9-eVLPs were added directly to the culture media supplemented with polybrene in each well. The transduced cells were then expanded into 6-well plate and sorted into single cell on FACS AriaIII sorter. A549 clones were then screened for their expression of the target gene by Western blot analysis. We used Western blot analysis to screen for KO clones. Three KO clones for STAT2 and four KO clones for IFIT1 were chosen for subsequent experiments. One KO clone and two partial KO clones for PAF1 were chosen for experiment in Figure 3A. The genomic DNA of the KO clones were extracted using PureLink Genomic DNA mini kit (Thermo Fisher Scientific) according to the manufaturer’s protocol. The targeted region of each gene was amplified by PCR, gel-purified, and subcloned into pUC57 plasmid by Gibson assembly. Between 15-20 plasmid clones were sequenced for each KO clones. Western blot analysis for IFIT1, STAT2, and PAF1 were performed for all the KO clones to confirm their expression profiles. For analysis in Fig. 3A-D and 3F, we used 3 STAT2-KO clones (C4, C12, F2), 4 IFIT1-KO clones (2B4, 2B3, 1A4, 1B3), and 3 PAF1-KO clones (3B (complete KO), 3F4 and 1B5 (partial KO)). For proteomic analysis (Fig. 3E), we used IFIT1-KO clone 2B3. For Fig. S8A, we used STAT2-KO clone C12 and PAF1-KO clone 3B. Two guide RNAs were used to knock out STAT2 and IFIT1, while one guide RNA was used for knocking out PAF1. The guide RNAs were designed using web tool provided by Synthego (https://www.synthego.com/crispr-design-tools/). The spacer sequences of the guide RNAs are provided in Table S1.

### Analysis of virus replication in human MDDC, M1, and M2

To assess viral replication in monocyte-derived dendritic cells, macrophages, cells were inoculated with the virus at a multiplicity of infection (MOI) of 5 for 2 hours. Subsequently, the cells were washed twice with RPMI 1640 and maintained in RPMI 1640 supplemented with 10% FBS for 24 hours. The percentage of cells expressing the DENV NS3 protein and the virus replication level (utilized as viral replication markers) were determined by flow cytometry. Cells were fixed with 4% formaldehyde, permeabilized with 0.5% saponin, and stained with an APC-conjugated anti-NS3 antibody (clone E1D8). Infectious viral particles released into the culture supernatant were quantified using a foci forming assay.

### Analysis of type-I IFN response in human MDDC, M1, and M2

To evaluate the induction of the type-I interferon response, we measured the intracellular levels of STAT1, STAT2, pSTAT1 (phosphorylated STAT1), and pSTAT2 (phosphorylated STAT2) as indicators of type-I interferon signaling activation. At 24 hours post-infection, cells were harvested, fixed with 4% formaldehyde, and permeabilized with 0.5% saponin. Intracellular staining was performed using rabbit monoclonal primary antibodies specific to the target proteins (Cell Signaling Technology), followed by anti-rabbit secondary antibodies (Cell Signaling Technology). Analysis was performed via flow cytometry, measuring STAT1 and STAT2 levels in both infected and uninfected cell populations to assess the type-I IFN response in both contexts.

### Analysis of human MDDC activation

To assess dendritic cell activation, we measured the expression of surface molecules, including HLA-class I, HLA-class II, HLA-E, CD80, CD83, CD86, and CD40. Staining was performed using fluorophore-conjugated monoclonal antibodies: Anti-HLA-class I, HLA-class II, and CD80 were obtained from BD; Anti-HLA-E, CD83, CD86, and CD40 were obtained from BioLegend. Cells were fixed with 4% formaldehyde and permeabilized with 0.5% saponin, followed by antibody staining at 4°C for 30 minutes. Results were analyzed using a flow cytometer.

### Cytokine and chemokine measurement

Cytokines and chemokines secreted by DENV-infected dendritic cells and macrophages were quantified in cell culture supernatants. Levels of IFN-α2, IFN-β, TNF-α, IL-12p70, IL-15, IL-18, CXCL-9, CXCL-10, CXCL-11, CXCL-8, CCL-3, and CCL-5 were measured using LEGENDplex™ assays. Data were acquired on BD-Fortessa, which differentiates bead populations and quantifies the PE signal intensity, allowing for the calculation of analyte concentrations based on a standard curve.

### Proteomic analysis

37,500 A549 cells were seeded in each well of 48-well plate and infected with DENV2 at MOI = 0.1 in RMPI1650 supplemented with 10% HI-FCS (R10) the next day for 2 hours. Cells were washed with 1xPBS twice before re-supplemented with fresh R10. The infected cells were maintained at 37°C, 5% carbon dioxide, with relative humidity between 90-95% for 3 days before harvest. At harvest, the plate was put on ice and then washed with 500 µl/well ice-cold 1XPBS, twice. Then, 6 wells of the cells were lysed with 100 µl RIPA buffer (50mM Tris-HCl pH7.5, 150mM NaCl, 1% NP-40, 0.5% sodium deoxycholate, 0.1% SDS, supplemented with 1x protease inhibitors). Protein concentration of lysate was determined using the BCA assay. For proteomic analysis, 50 μg of total protein was subjected to methanol–chloroform precipitation. The resulting protein pellets were washed with methanol, disrupted by sonication, and resuspended in 50 mM Tris buffer (pH 8.0) at a concentration of 1 mg/mL. Proteins were reduced with 5 mM Tris(2-carboxyethyl)phosphine (TCEP) and alkylated with 15 mM chloroacetamide (CAA) for 45 min at room temperature. Proteolytic digestion was performed overnight at 37 °C with sequencing-grade trypsin (Promega) at a 1:100 w/w enzyme-to-protein ratio. Peptides were subsequently desalted using triple-stacked SDB-XC StageTips before LC-MS/MS analysis.

Dried peptides were reconstituted in 0.1% formic acid (FA) in LC-MS-grade water and transferred into low-adsorption vials (Supelco, Merck). Samples were analyzed on a Dionex Ultimate 3000 nRSLC system coupled to a Q Exactive™ HF Orbitrap mass spectrometer (Thermo Fisher Scientific) equipped with an EASY-Spray™ ion source. Peptides were initially captured on a PepMap™ Neo C18 trap column (5 μm, 300 μm × 5 mm) and separated on an EASY-Spray™ Acclaim PepMap C18 analytical column (50 cm × 75 μm). Chromatographic separation was achieved using a 125-min linear gradient ranging from 0 to 85% solvent B (0.1% FA in acetonitrile) at a flow rate of 300 nL/min. Data were acquired in data-dependent acquisition (DDA) mode using a TopN15 method. Full MS scans were recorded over an *m/z* range of 375-1300 at a resolution of 120,000 (FWHM at *m/z* 200), with an AGC target of 3×10^6^and maximum injection time of 25 ms. MS/MS spectra were acquired at a resolution of 15,000 (FWHM at *m/z* 200), with an AGC target values of 5×10^4^, and a maximum ion injection time of 35 ms. The 15 most intense precursor ions with charge ≥2 were selected from each MS1 scan using a 1.6 *m/z* isolation window and fragmented by higher-energy collisional dissociation (HCD) with a normalized collision energy (NCE) of 28 on the scan range of 200 to 2000 *m/z*.

Raw MS data were processed using FragPipe (v.22.0), with MSFragger (v.4.0)(*71*). The built-in “LFQ-MBR” workflow was applied with precursor and fragment mass tolerances set to 40 ppm and 10 ppm, respectively. Searches were conducted against the reviewed human proteome database (UniProt ID: UP000005640_9606, downloaded December 2024), containing 20,638 protein sequences along with their corresponding decoy sequences and common contaminants. Peptide-spectrum matches and protein identifications were filtered to achieve a false discovery rate (FDR) below 1% using a target-decoy strategy, with a minimum peptide length of seven amino acids. Carbamidomethylation of cysteine residues was specified as a fixed modification, while methionine oxidation and protein N-terminal acetylation were included as variable modifications, allowing up to three modifications per peptide. Strict trypsin specificity was applied for digestion. Label-free quantification (LFQ) was performed using IonQuant (v.1.10.27)(*72*). Downstream data processing was carried out in Perseus (v.2.0.11.0)(*73*), and graphical visualization was generated using GraphPad. The list of ISGs used for analysis was based on previous report(*74*).

### Protein structure analysis

Contact analysis between NS5 and STAT2 was performed using ChimeraX version 1.6.1 using center-center distance less than or equal to 5Å (*75*).

### Principal component analysis and K-mean clustering

To reduce the dimensionality of the cytokine and phenotypic data while preserving variance, Principal Component Analysis (PCA) was performed using the scikit-learn library. The dataset, comprising multiple biological features (e.g., cytokine levels such as IFNa2 and CXCL9 across different cell types like M1 and MDDC), was first standardized using a StandardScaler to ensure each feature had a mean of zero and unit variance. PCA was then applied with two target components (PC1 and PC2). The explained variance ratio for the first two components was calculated to assess the proportion of information retained. To identify distinct biological groupings based on the integrated cytokine and phenotype profiles, K-means clustering was applied to the standardized feature matrix. The number of clusters (K) was defined based on the types of mutations (K=5) in the DENV2 panel. The algorithm was initialized with a fixed random state of 42 to ensure reproducibility and used 10 initializations (n_init=10) to find the most stable cluster centroids. Clusters were visualized by projecting the results onto the two-dimensional PCA space, using distinct markers and colors for each identified cluster (Fig. 4A).

### Statistical analysis of immune profiles from primary human immune cells across DENV2 panel

Data stratified by donor, cell type, and infection condition were normalized using z-score normalization via the StandardScaler function in the scikit-learn library. To determine whether viral mutations significantly altered cytokine/chemokine expression or phenotypic profiles, statistical comparisons were performed using a one-way Analysis of Variance (ANOVA) for each cytokine across all viral groups (e.g., WT, PDK53, and specific mutants). Data were analyzed independently for each cell type (M1, M2, and MDDC). For cytokines/chemokines or phenotypic markers demonstrating a statistically significant difference by ANOVA (α = 0.05), Tukey’s Honestly Significant Difference (HSD) post-hoc test was employed to identify specific pairwise differences between viruses while controlling for the family-wise error rate. Statistical computations were implemented in Python using the scipy.stats and statsmodels libraries. P-value matrices were generated from the Tukey HSD results and transformed to a scale to represent statistical significance. These results were visualized as heatmaps using the seaborn and matplotlib packages. For infection markers (Fig. S9H-J), the data were not normalized before statistical analysis.

### Statistical analysis of phenotypes and cytokines/chemokines of mock vs. DENV infected

To evaluate the impact of viral infection on cytokine induction, statistical comparisons were performed between infected and mock-treated cells for each cytokine across three primary cell types (M1, M2, and MDDC). For each virus-cell-gene combination, an independent two-sample t-test was used to determine if the mean cytokine levels in the infected group differed significantly from the mock group. The data were visualized using bar plots generated with the seaborn and matplotlib libraries in Python. Error bars represent the standard error of the mean (SEM).

## Acknowledgments

The authors would like to thank Prida Malasit and Prasert Auewarakul for advice, Chunya Puttikhunt and Watchara Kasinrerk for sharing the anti-NS1 antibody used in this work.

## Funding

The work reported here was funded by the grants from Newton-NSTDA-MRC program in infectious diseases (P-18-50538 and GB-GOV-13-FUND-Newton-MR_R020957_1 to B.S. and J.M., respectively), from PMU-B (B16F650039 to B.S.) and from National Vaccine Institute (2567.1/8 to W.D.).

## Author contributions

Conceptualization: B.S, J.M.,W.D. ; Methodology: B.S, J.M.,W.D., A.S., W.C., P.P., N.P., G.S.; Experiments and Data Collections: B.S, W.D., A.S., W.C., P.P., Nittaya S., R.N., N.P., T.R., S.O., B.T.; Analysis: B.S, J.M., W.D., A.S., W.C., P.P., N.P.; Supervision: B.S, J.M.,W.D.; Writing—original draft: B.S., W.D., A.S., W.C., P.P., N.P.; Writing—review & editing: B.S, J.M.,W.D., G.S., Nopporn S., P.K.

## Competing interests

B.S., W.D., J.M., A.S., P.P., W.C., N.S., R.N., and S.O. have filed patent applications relating to this work.

## Data, code, and materials availability

The DMS data of DENV2 library infection in A549 +/-INF-α2 were deposited in MaveDB database (*76*) under accession number 93b955df-d883-458b-85a6-e6111dc87b2c. The raw deep sequencing data have been deposited in the Sequence Read Archive under BioProject accession number PRJNA1464240. The mass spectrometry proteomics data have been deposited to the ProteomeXchange Consortium via the PRIDE partner repository with the dataset identifier PXD077464.

## Supplementary Materials

**Table S1:**
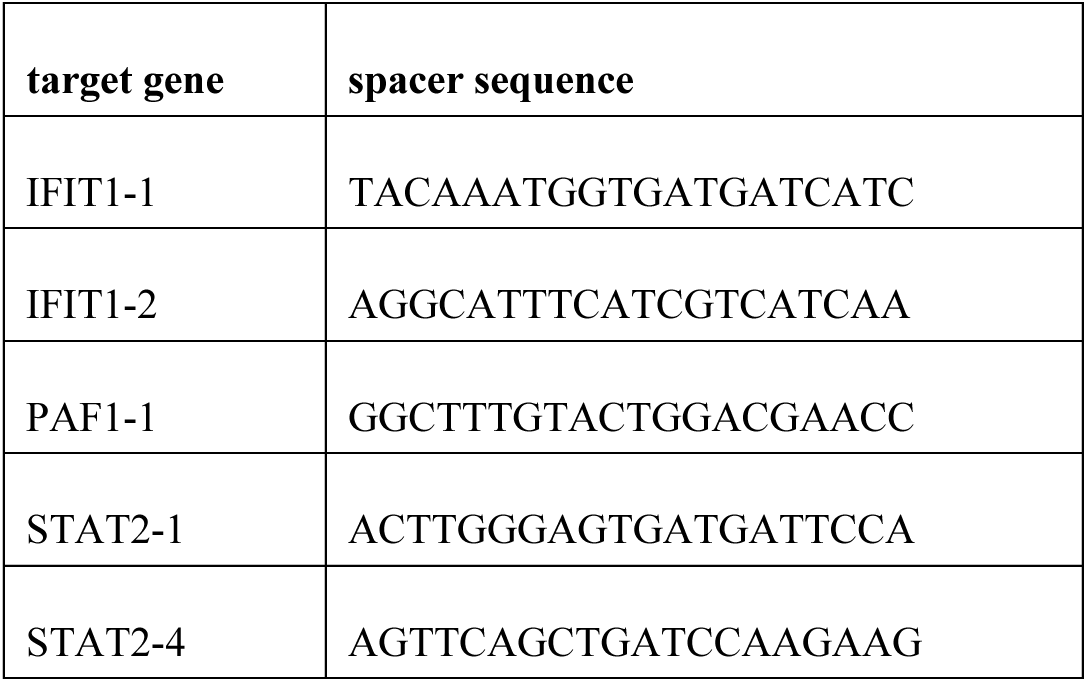
Spacer sequences of the guide RNAs used in this study.

**Fig. S1:**
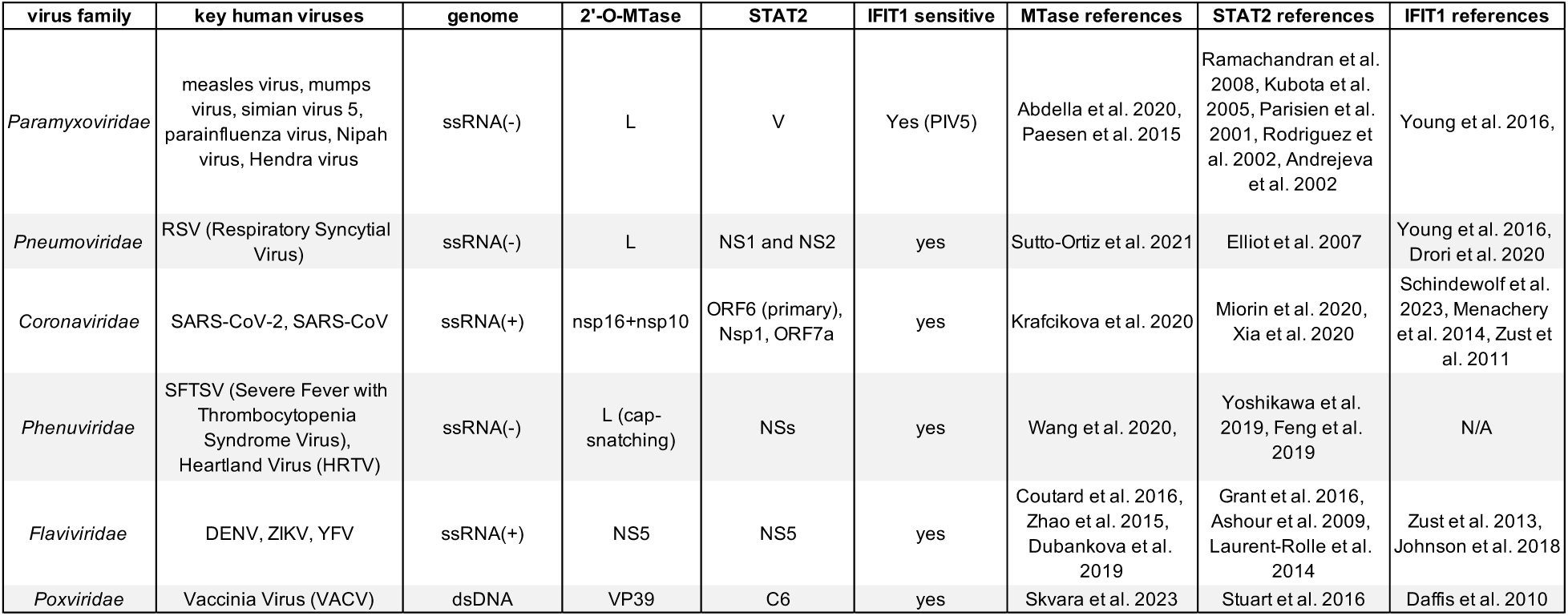
Human viruses with viral 2’-O-methyltransferase and STAT2-targeting proteins. The sensitivity to IFIT1 when their viral 2’-O-methylation has been compromised was demonstrated for most of the viruses shown here. The references are listed in the main text.

**Fig. S2:**
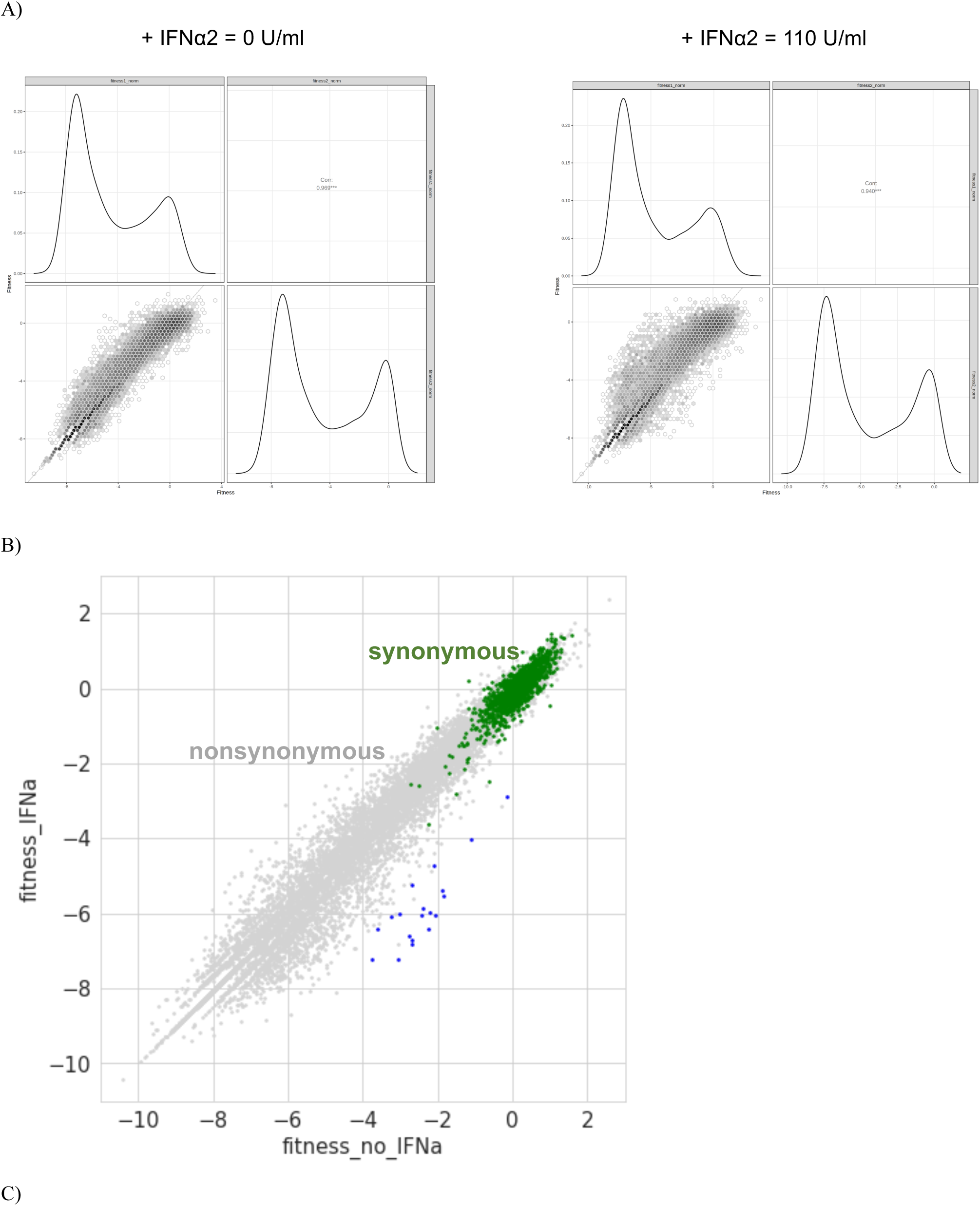
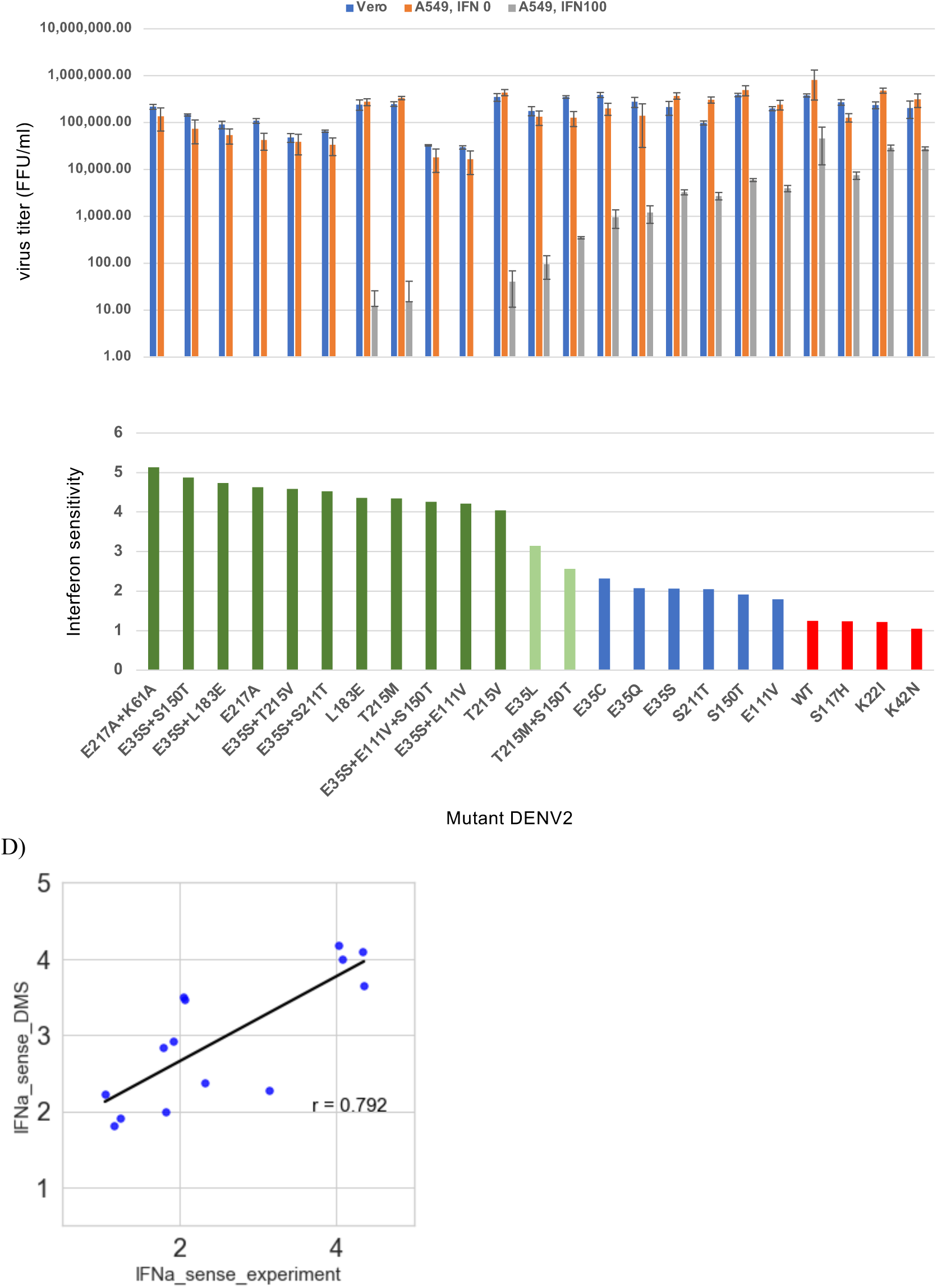
Additional DMS and validation data from Fig. 1A-C. A) Correlation of fitness scores from two replicates of DMS under the conditions with and without IFN-α2. B) Scatter plots showing the correlation of mutant fitness scores under the conditions with and without IFN-α2. The blue dots represent the mutants that have IFN sensitivity above 2.5 and DMS fitness scores in Vero cells higher than -3. (C) The infectious titers (top) and the interferon sensitivity (bottom) of the selected mutants in Vero and A549 +/- IFN-α2. D) Correlation between the experimental measurement of IFN sensitivity of single amino-acid mutants in C) with the corresponding IFN sensitivity calculated from the DMS fitness scores.

**Fig. S3:**
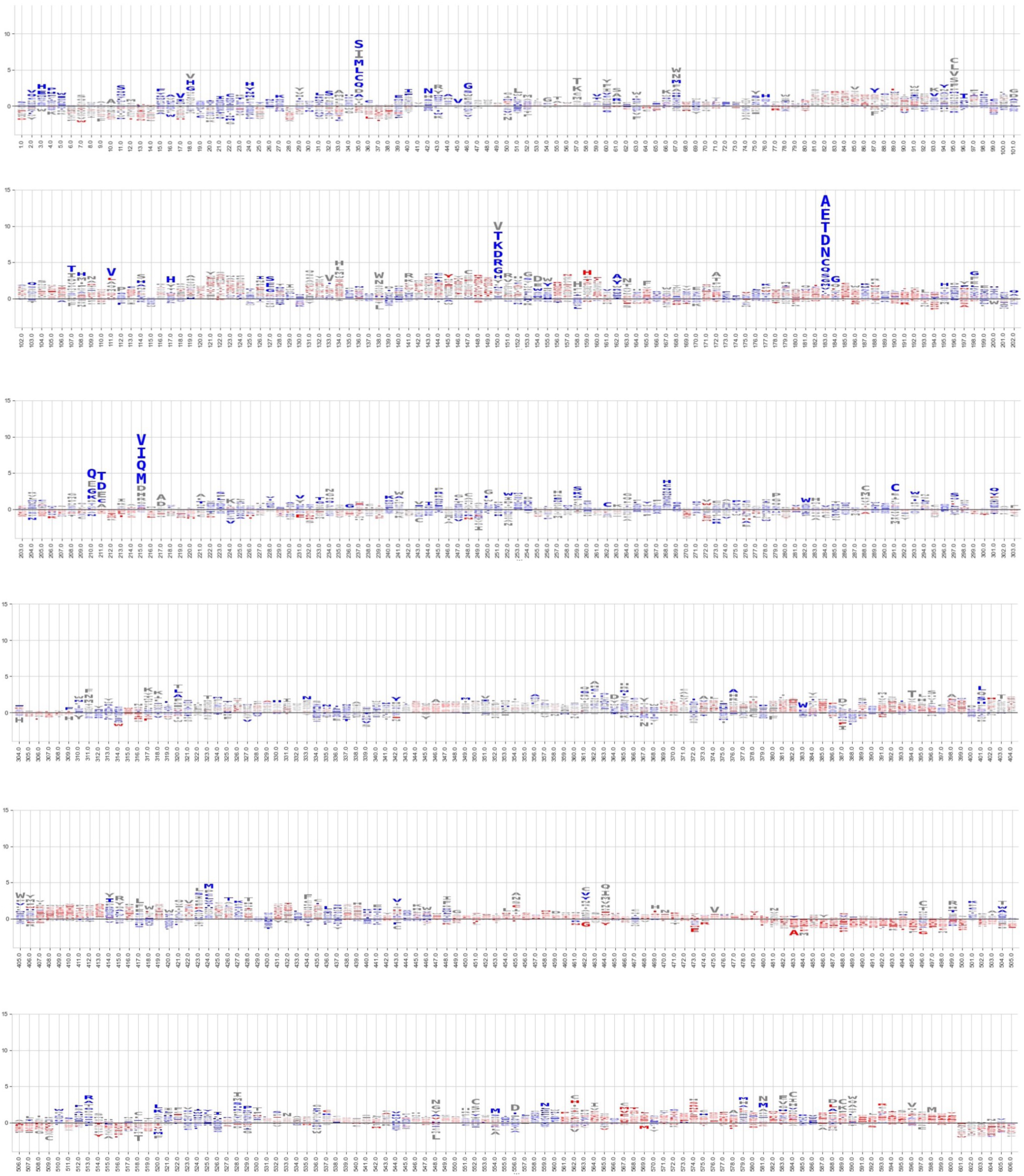
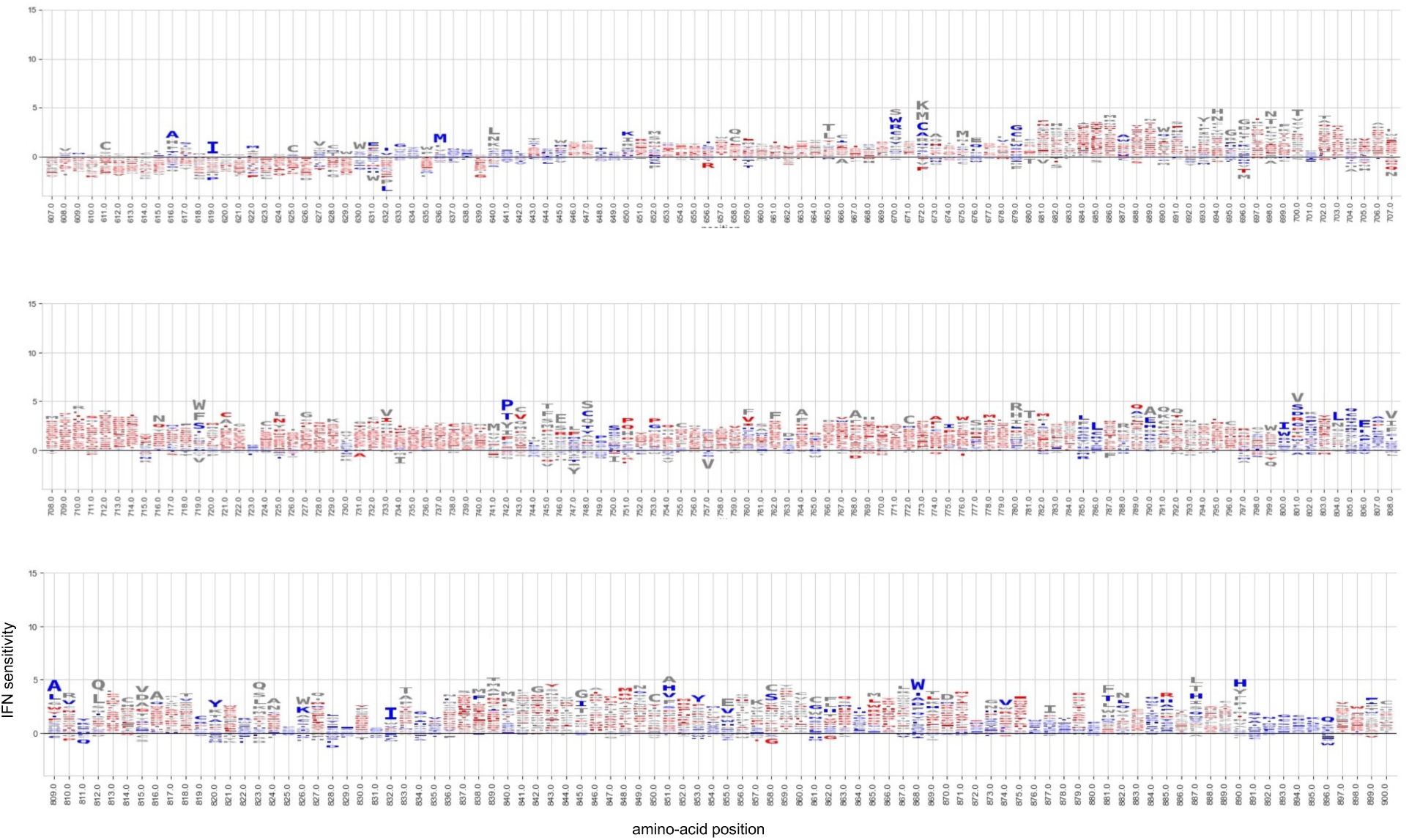
WebLogo plot of the IFN sensitivity of single amino-acid mutations across the length of DENV NS5. The size of the letter is proportional to the IFN sensitivity for each mutant.

**Fig. S4:**
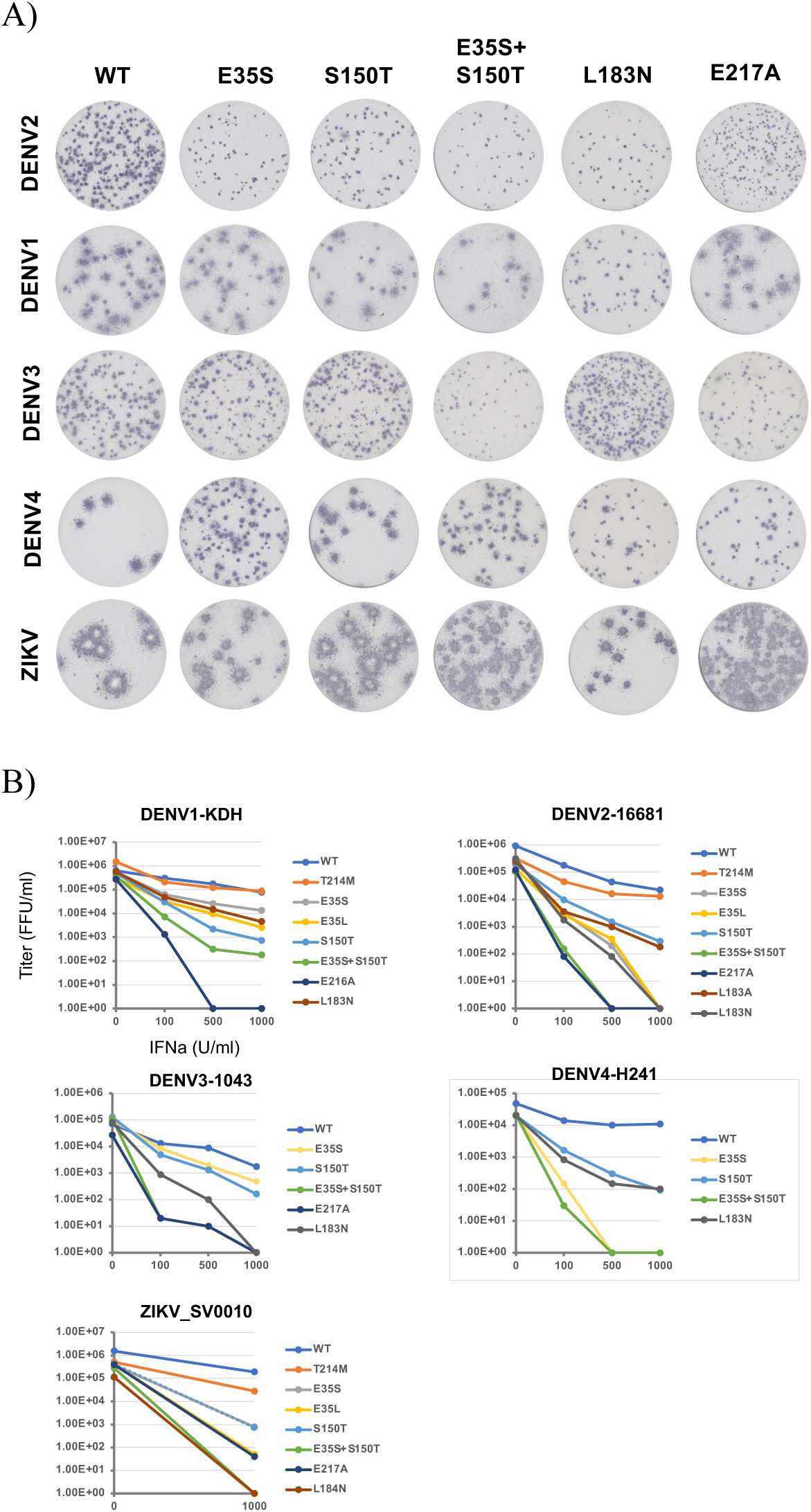
Additional data from Figure 1D on the interferon sensitivity of DENV1-4 and ZIKV with MTase mutations. A) Foci images of the DENV1-4 and ZIKV with the MTase mutations used in this study. B) Dose-response curves of the mutant viruses. The horizontal axes correspond to the IFN-α2 dosage. The vertical axes correspond to the virus titer measured at 2 dpi. A549 was infected with a virus at MOI = 0.1. The cells were pre-treated with IFN-α2 at 4 hours pre infection.

**Fig. S5:**
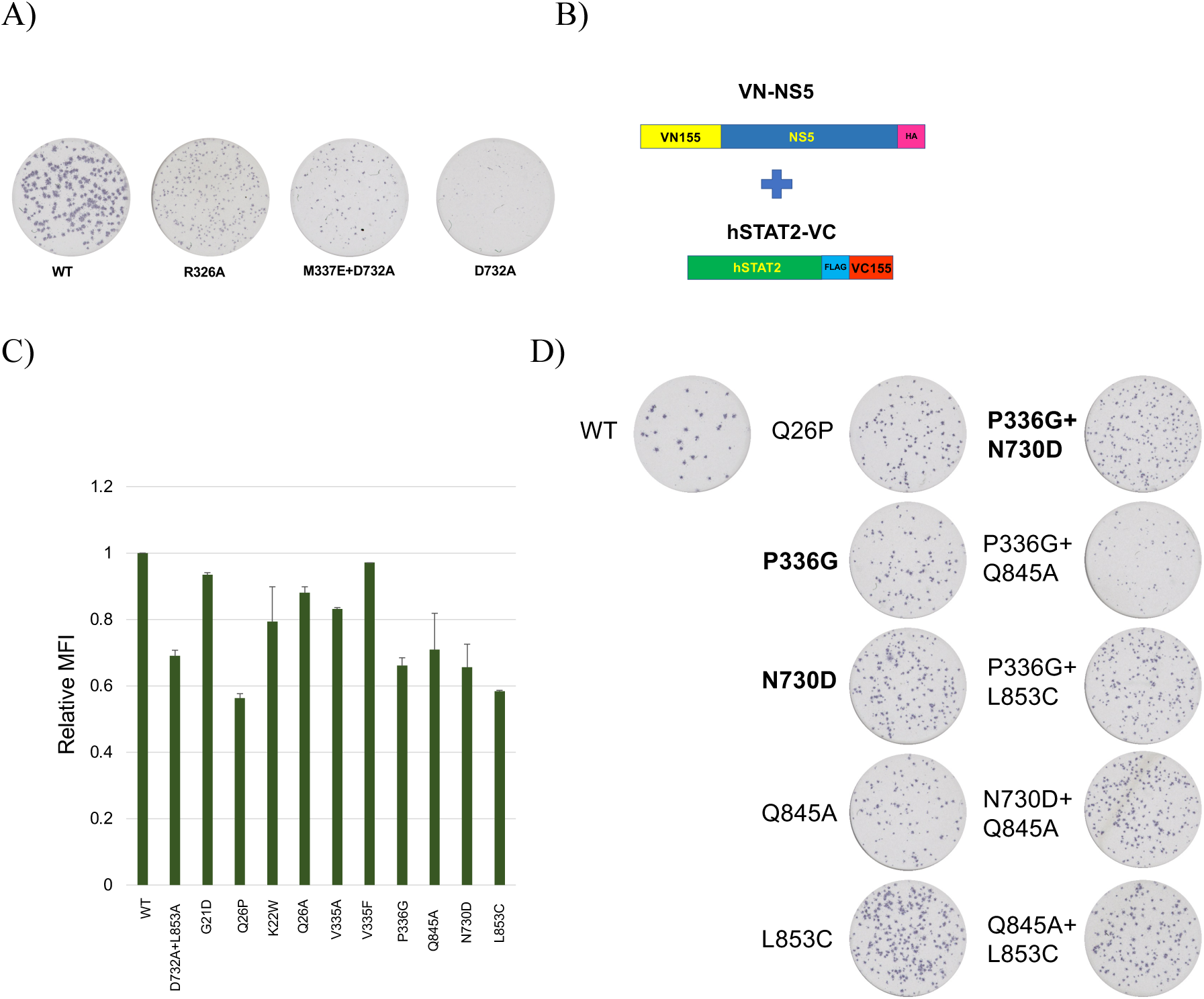
Additional data on the identification of NS5 mutations that disrupt STAT2 targeting by BiFC. A) Foci images of the DENV2 with ZIKV-equivalent STAT2-targeting mutations. B) Design of bimolecular fluorescence complementation (BiFC) assay for NS5-STAT2 interaction analysis in cells. C) Flow cytometry BiFC readouts of selected NS5 mutants with human STAT2. D) Foci images of DENV2 with single and combined STAT2-targeting mutations.

**Fig. S6:**
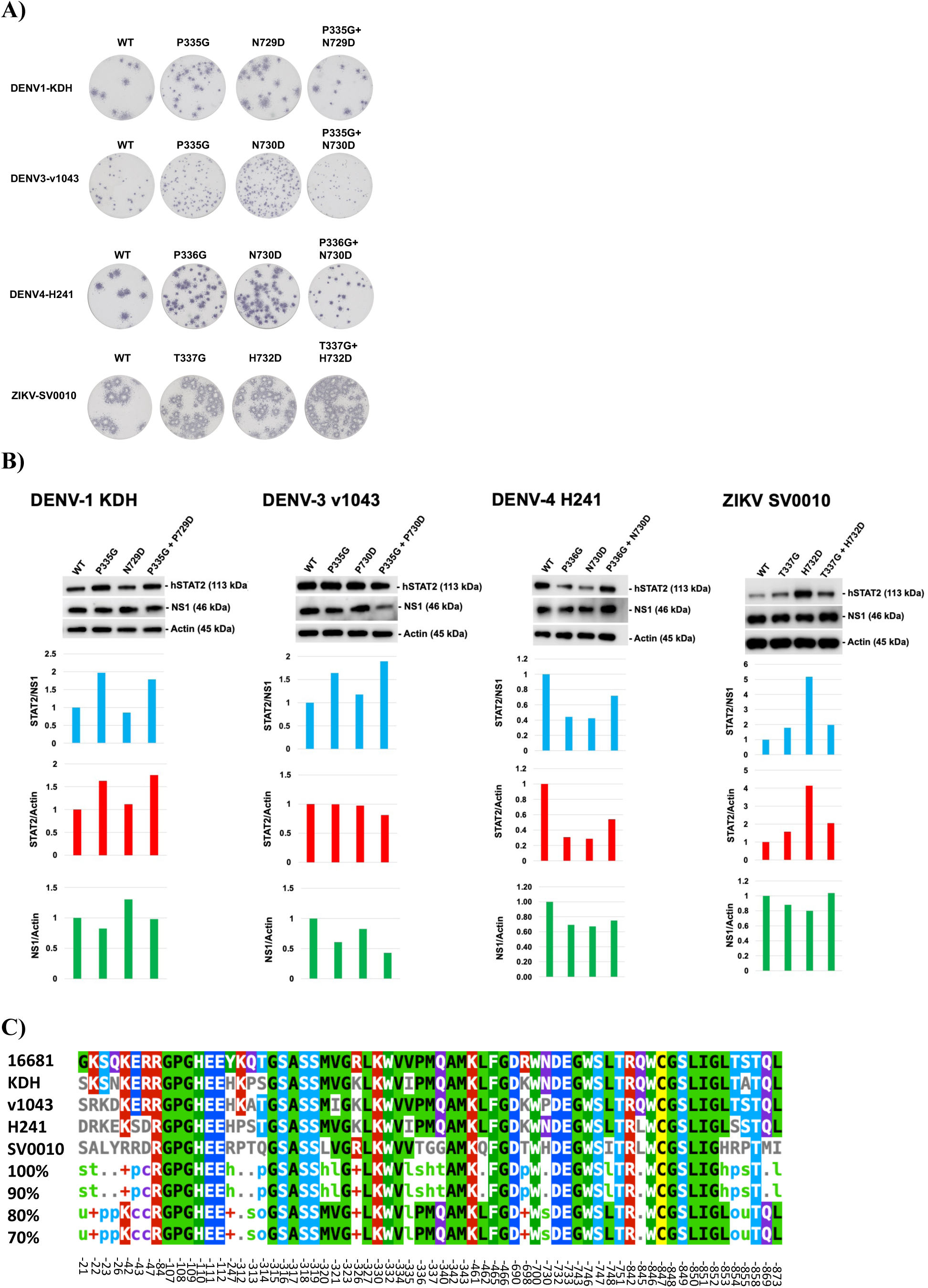
Characterization of DENV2 STAT2-targeting mutations in DENV1-4 and ZIKV backgrounds. A) Foci images of DENV1-4 and ZIKV with P336G or N730D (DENV2 mutations). B) STAT2 degradation in A549 cells infected by DENV1-4 and ZIKV with P336G or N730D mutations. C) Multi-sequence alignment (MSA) of STAT2-interating residues across DENV1-4 (KDH, 16681, v1043, and H241, respectively) and ZIKV (SV0010) used in A) and B). MSA was performed with full-length NS5 sequences. The STAT2 interacting residues (based on the analysis performed in Fig. 2) were then extracted from the aligned sequences and displayed using MView (78).

**Fig. S7:**
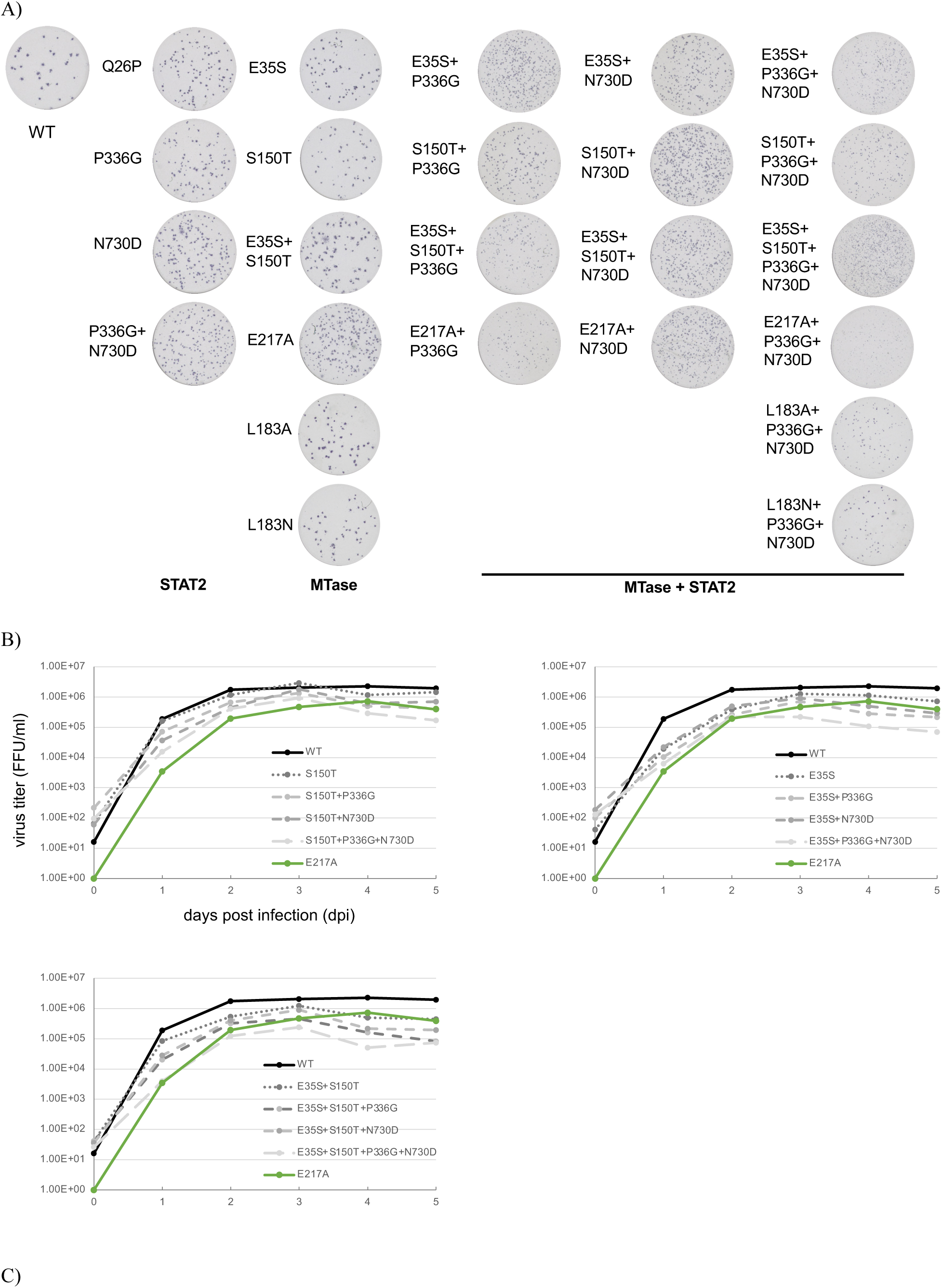
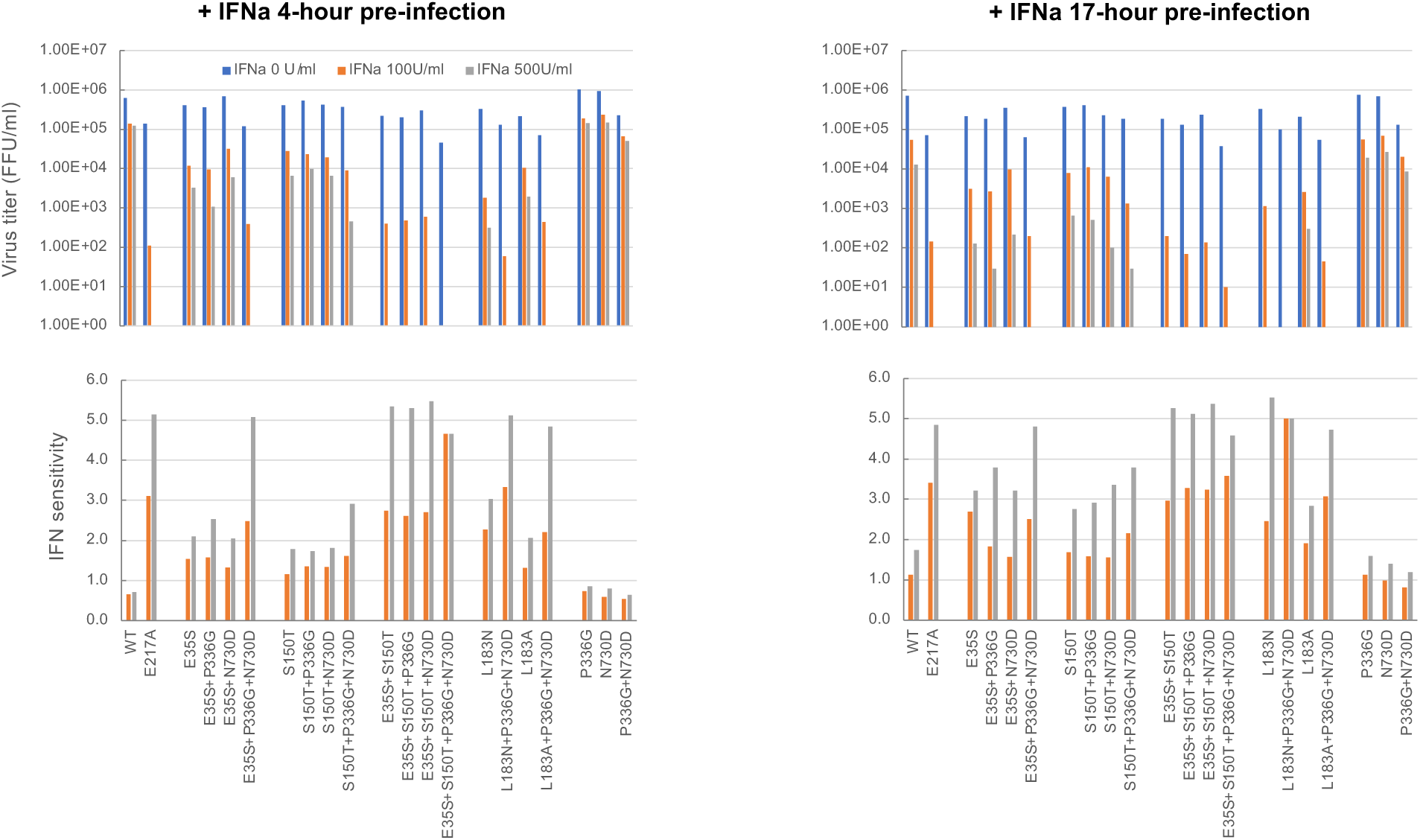
Characterization of basal replicative fitness and IFN sensitivity of DENV2 with combined MTase and STAT2-targeting mutations in Vero cells. A) Foci images of the mutant DENV2. B) Multi-step replication kinetics of the mutant viruses in Vero cells. Vero cells were infected at MOI = 0.1. C) The infectious titers (top) and the interferon sensitivity (bottom) of the selected mutants in A549 +/- IFN-α2 at 100U/ml or 500U/ml and pre-treated at 4 and 17 hours pre-infection. A549 cells were infected with a virus at MOI = 0.1. The virus titers were measured 2 days post infection.

**Fig. S8:**
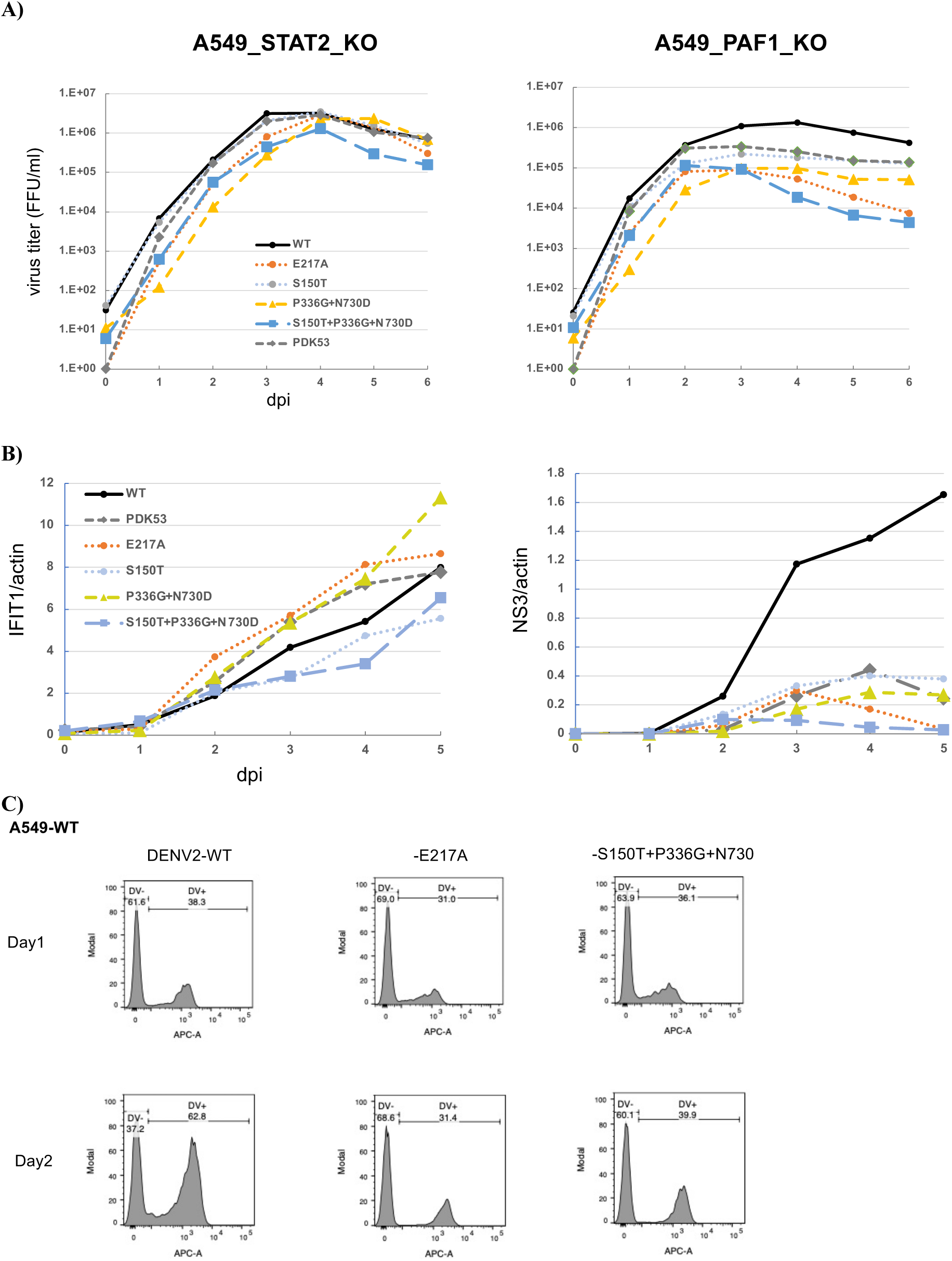
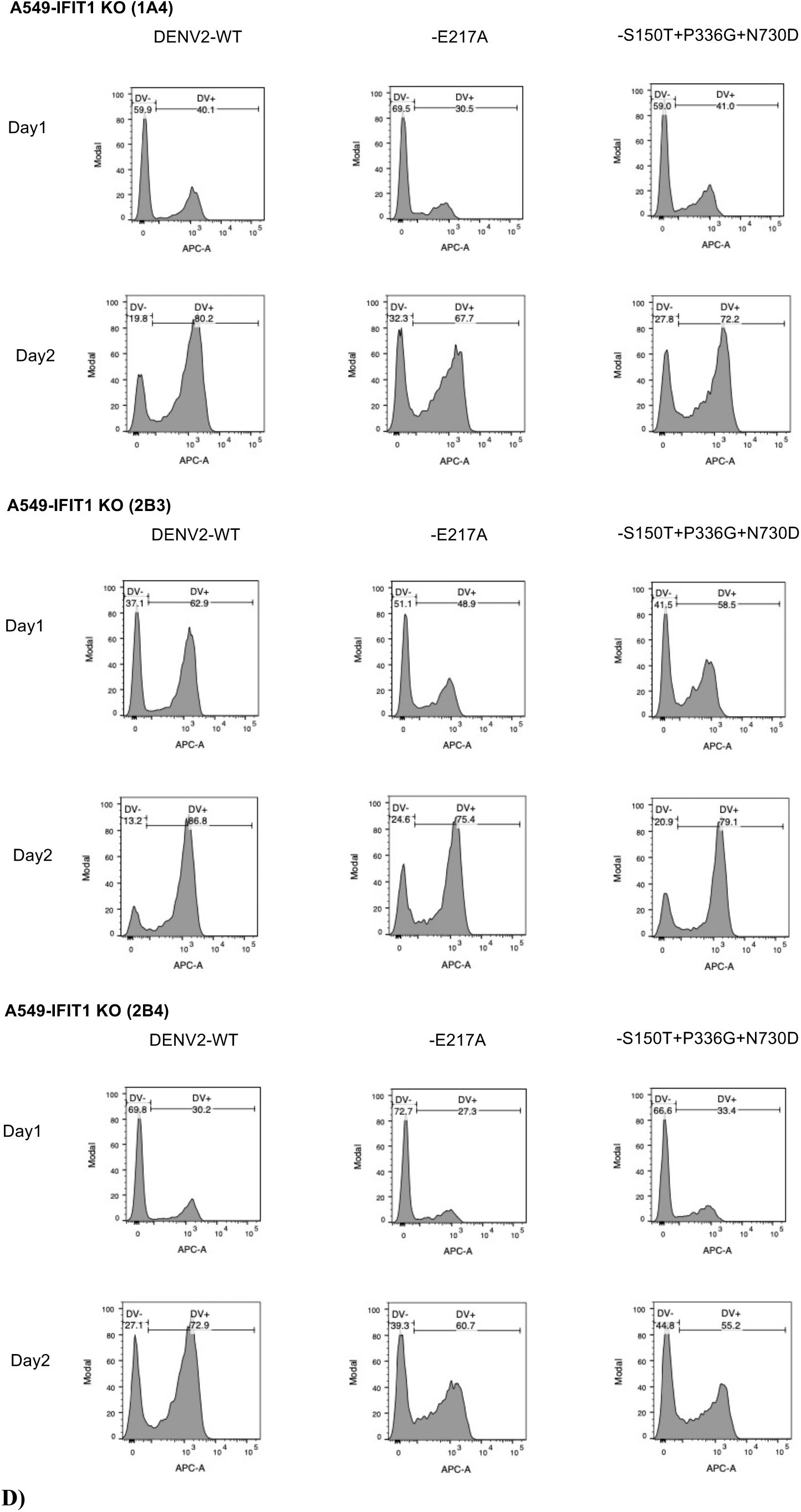
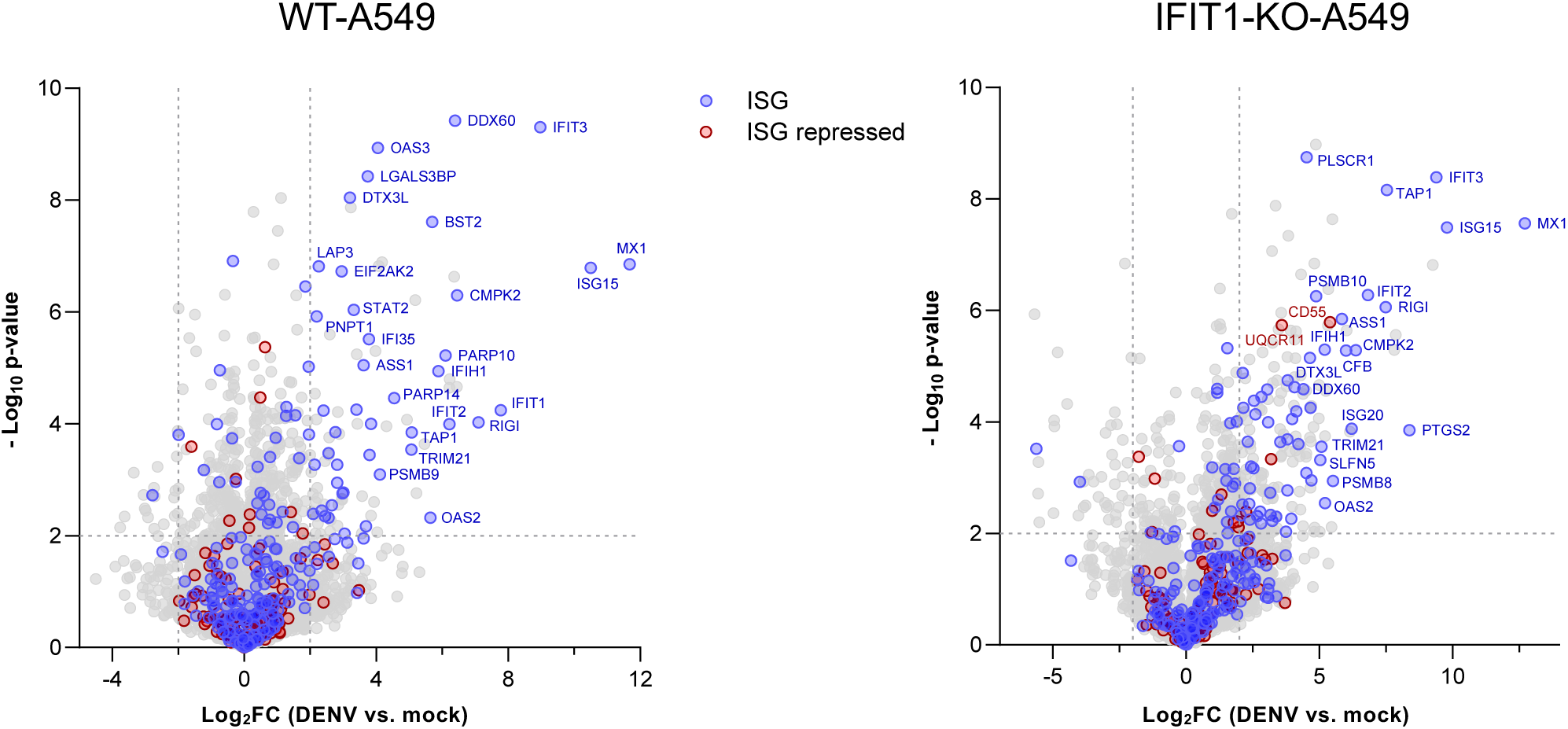
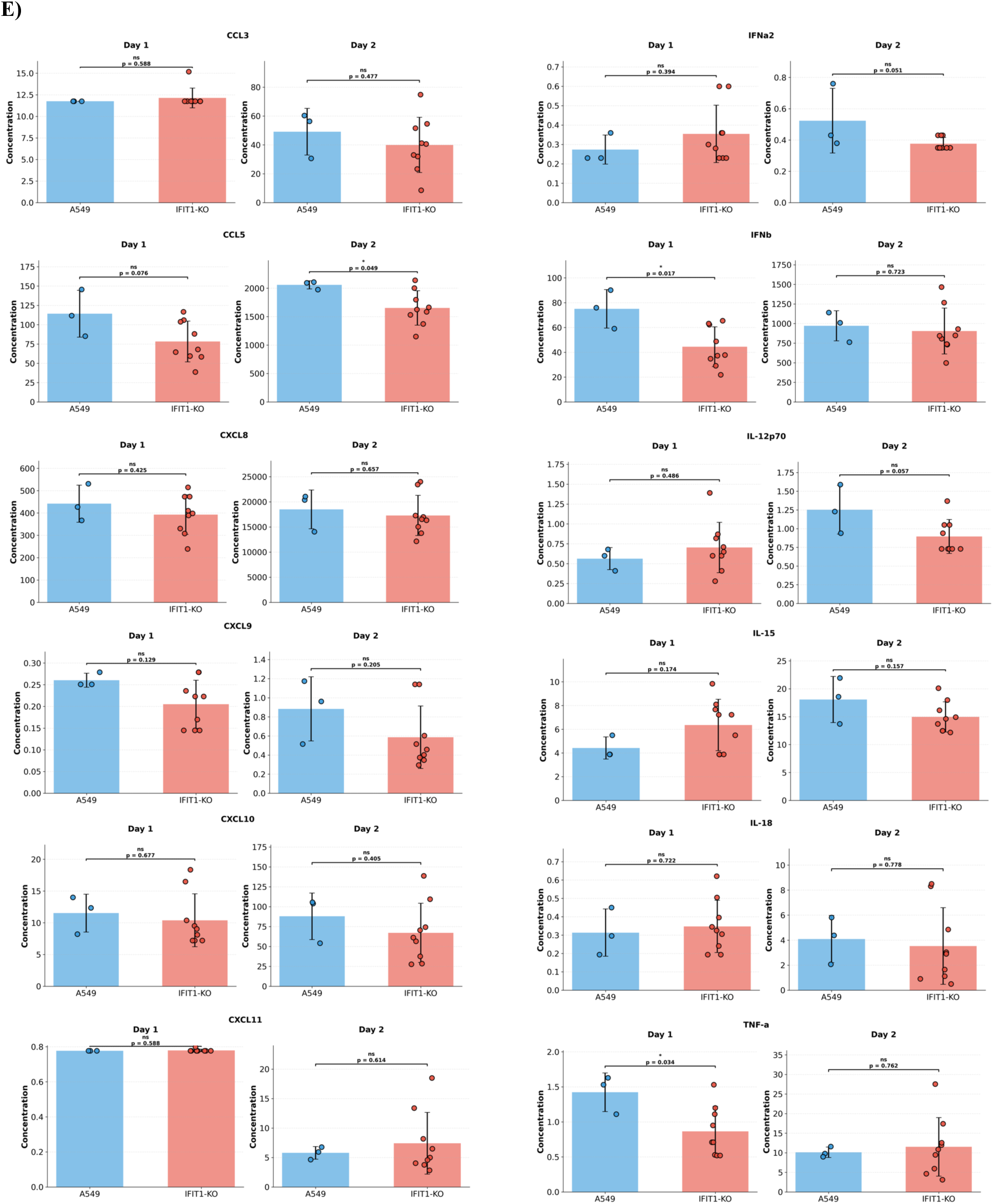
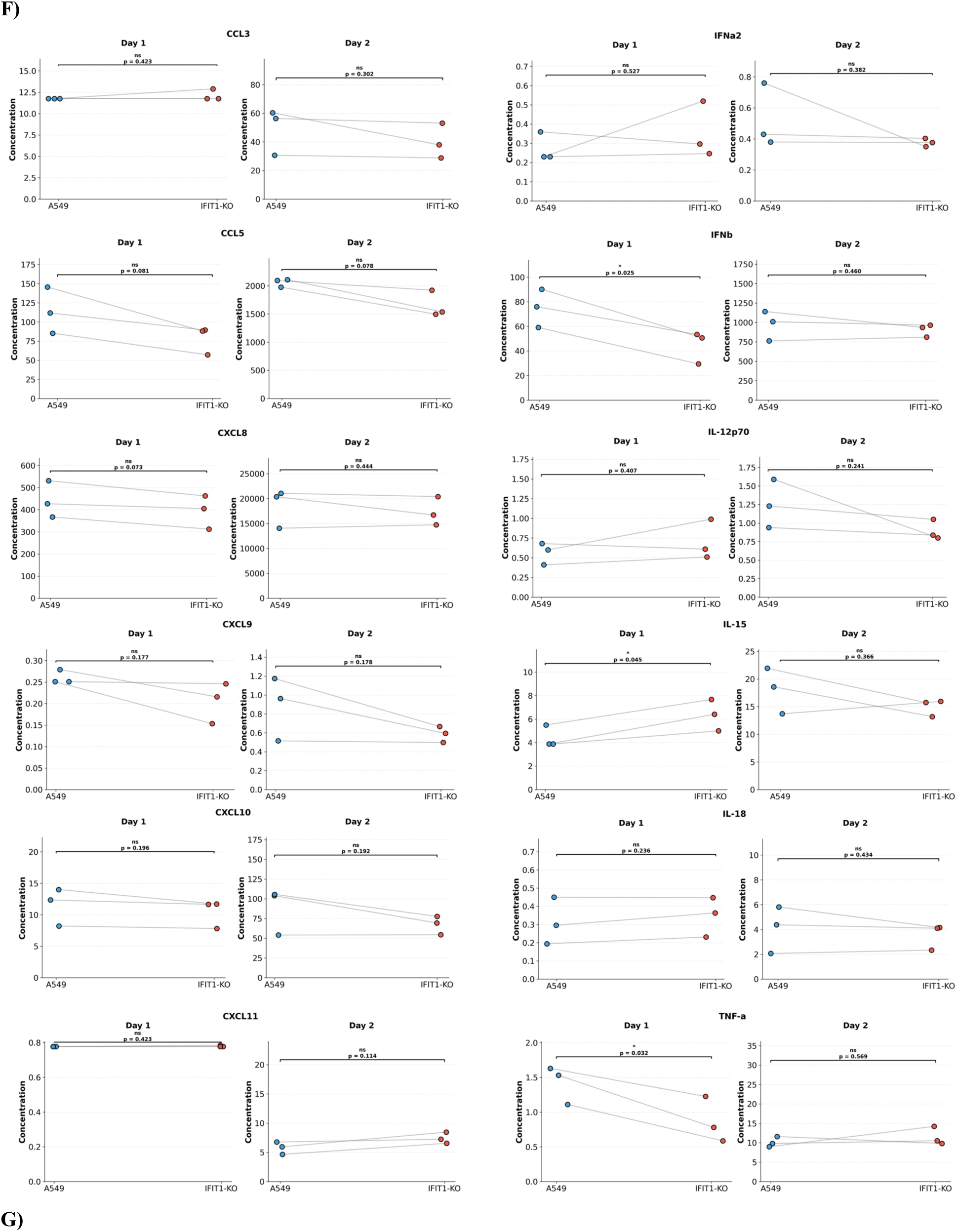
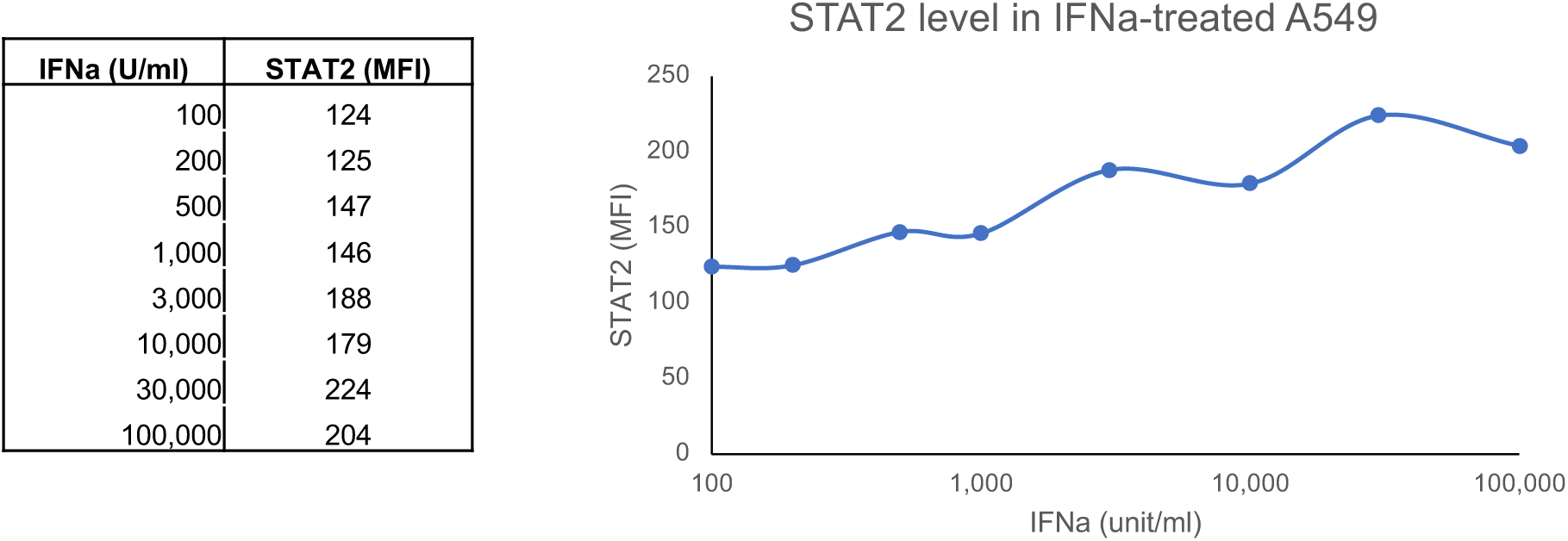
Additional analyses to characterize IFIT1-STAT2 signaling relationship. A) Multi-step replication kinetics of the mutant DENV2 in STAT2-KO and PAF1-KO A549 cells. A549 cells were infected with DENV2 at MOI = 0.1. B) Western-blot analysis of IFIT1 and dengue NS3 expression in wild-type A549 cells during multi-step replication kinetics analysis in A). C) Flow cytometry of DENV2 infection by anti-NS3 in WT and IFIT1-KO A549 cells. D) Volcano plots illustrating the distribution of quantified proteins. The x-axis represents the fold change (LFC), and the y-axis represents the statistical significance as (-value). Proteins with a fold change > 2 and a p-value < 0.01 were considered significantly differentially expressed. Selected ISGs upregulated during DENV infections in WT- and IFIT1-KO A549 are labeled by their gene symbols. Aggregate data of DENV2-E217A and DENV2-S150T+P336G+N730D were used to represent the data of infected cells. E-F) Comparison between the cytokine concentrations (pg/ml) secreted from WT-A549 and those from IFIT1-KO-A549 infected with one of the three viruses, DENV2-WT, -E217A, or S150T+P336G+N730D (MOI = 1) at 1 and 2 dpi. The data points in the top bar plots (E) from each virus were aggregated by cell types. The bottom paired line plots (F) were matched by virus, with the data points in the IFIT1-KO being an average of the measurement from three IFIT1-KO clones. Statistical significance is indicated by brackets above the data sets. Exact values are provided for each comparison (* ; ns, not significant). Statistical analysis was performed using an unpaired Student’s t-test for the top bar plots and a paired t-test for the bottom line plots. G) Dose-response analysis of IFN-α2 and STAT2 level in A549 cells. A549 cells were treated with IFN-α2 48 hours before measurement by flow cytometry.

**Fig. S9:**
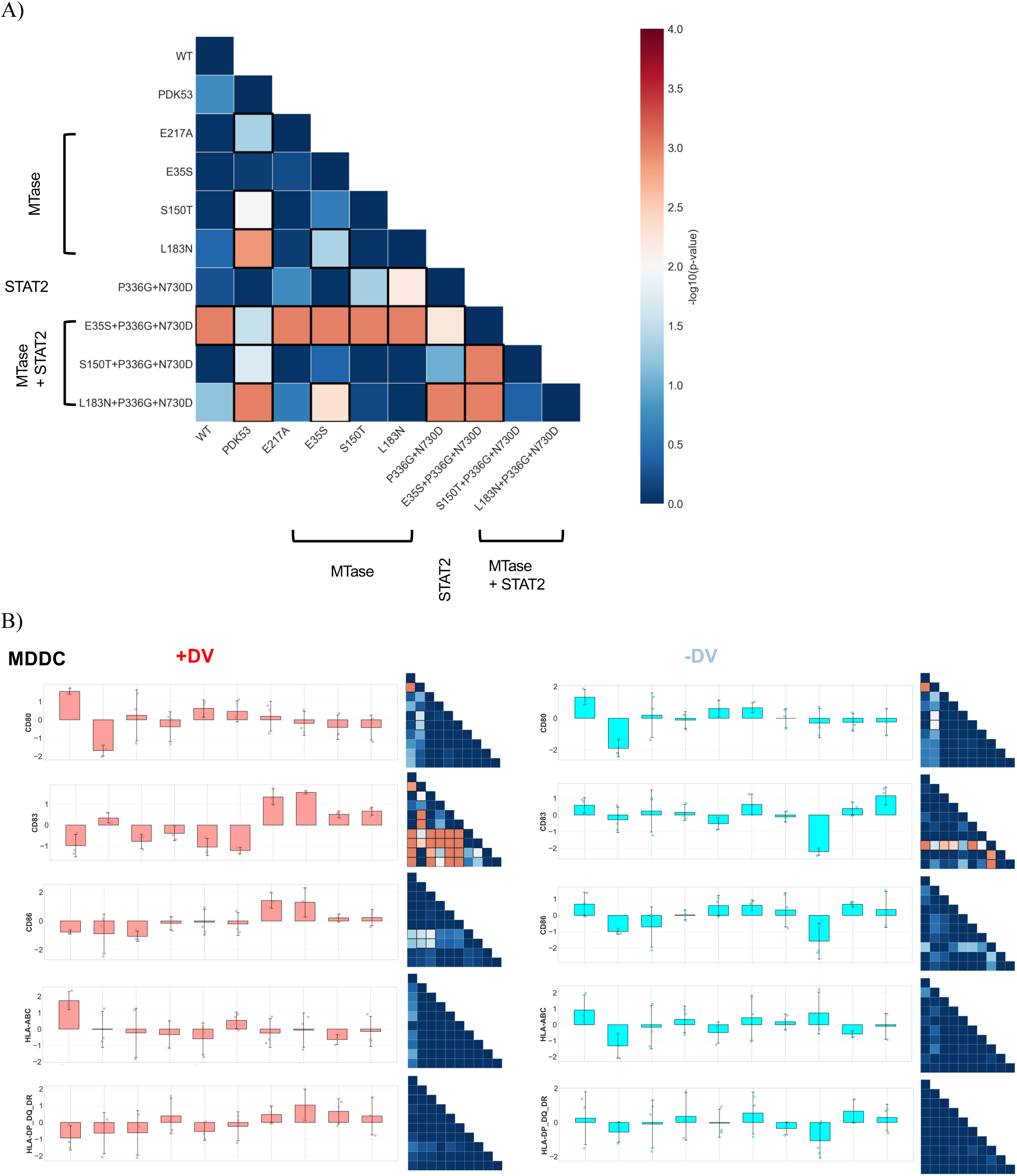
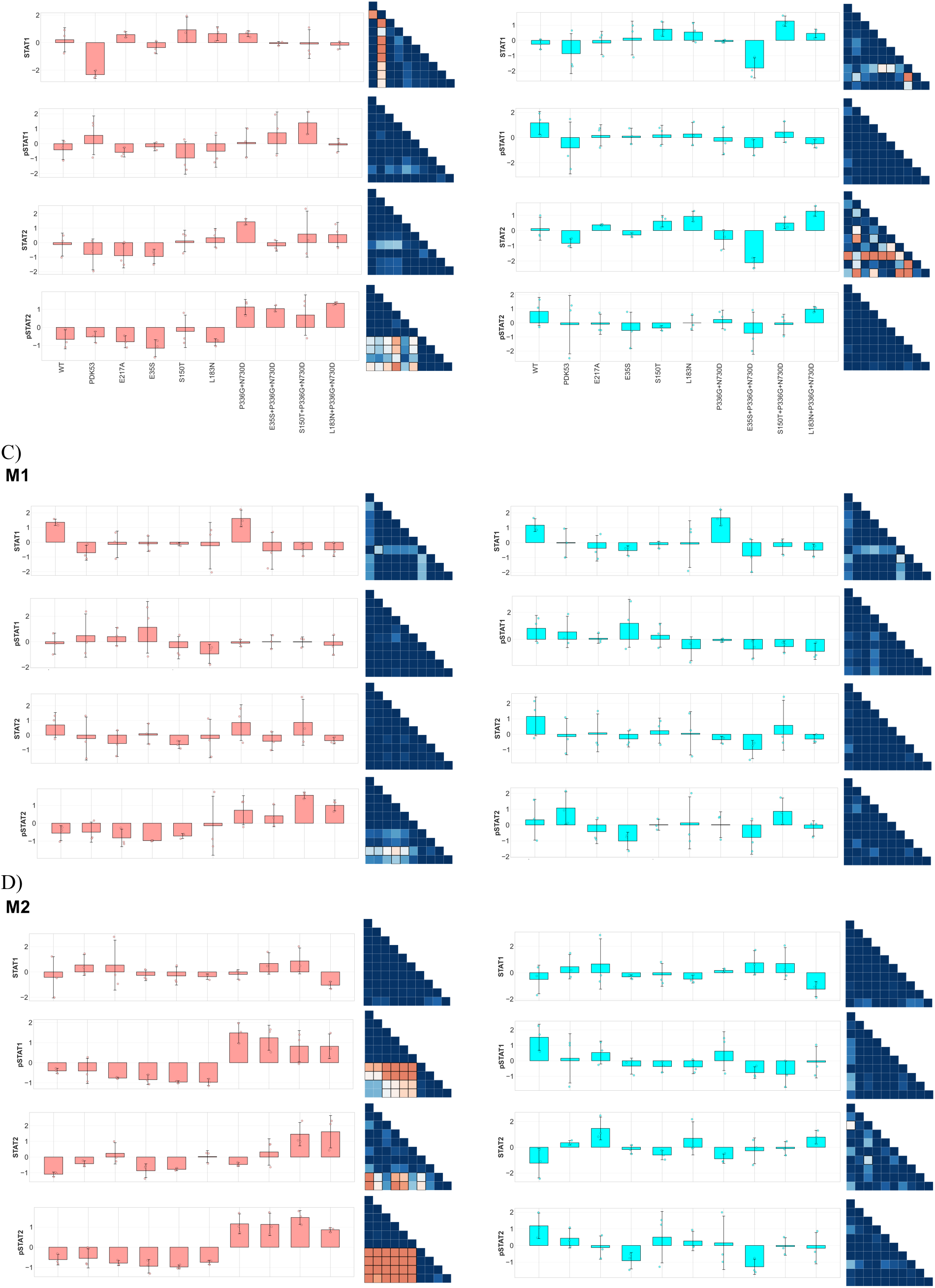
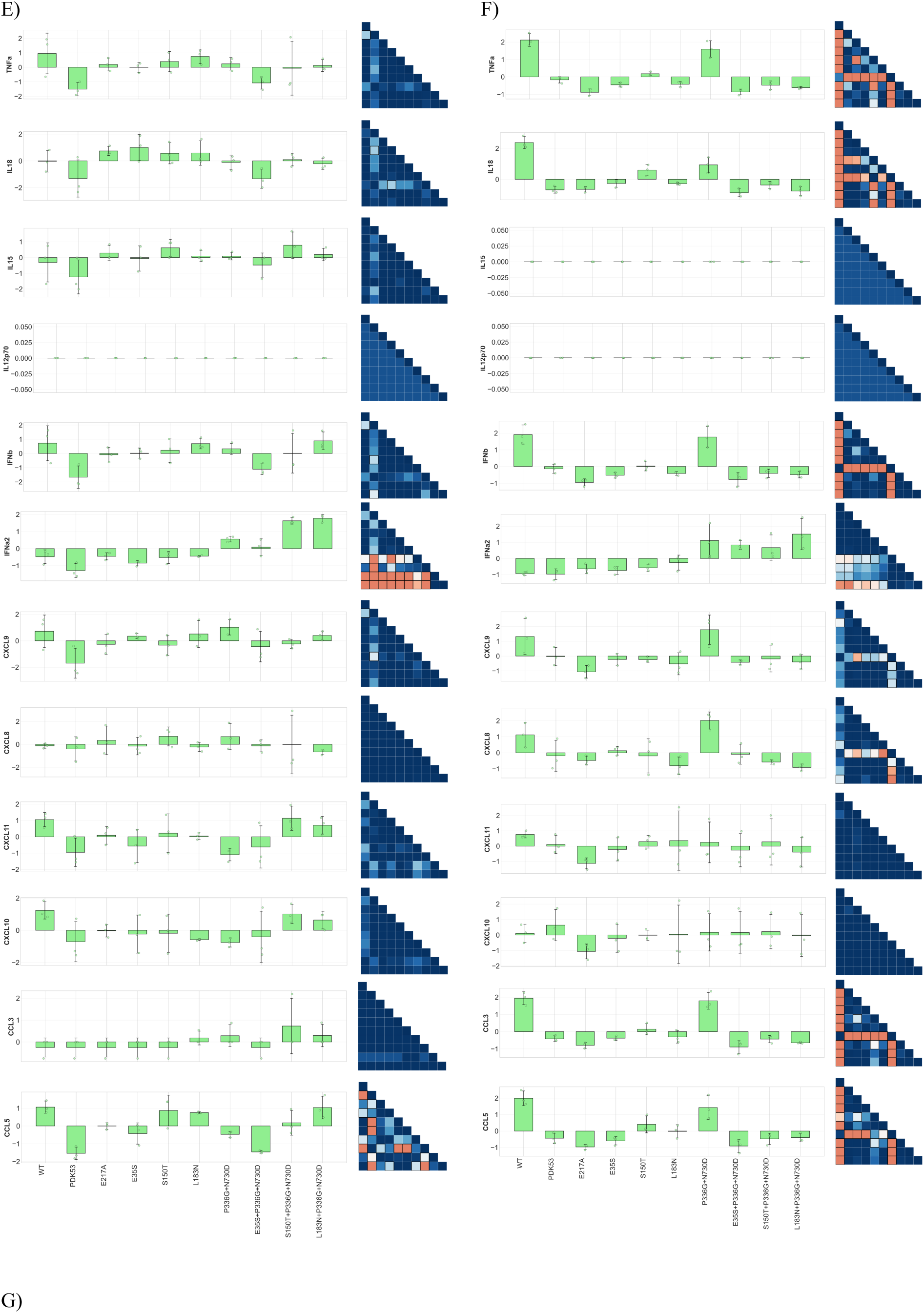
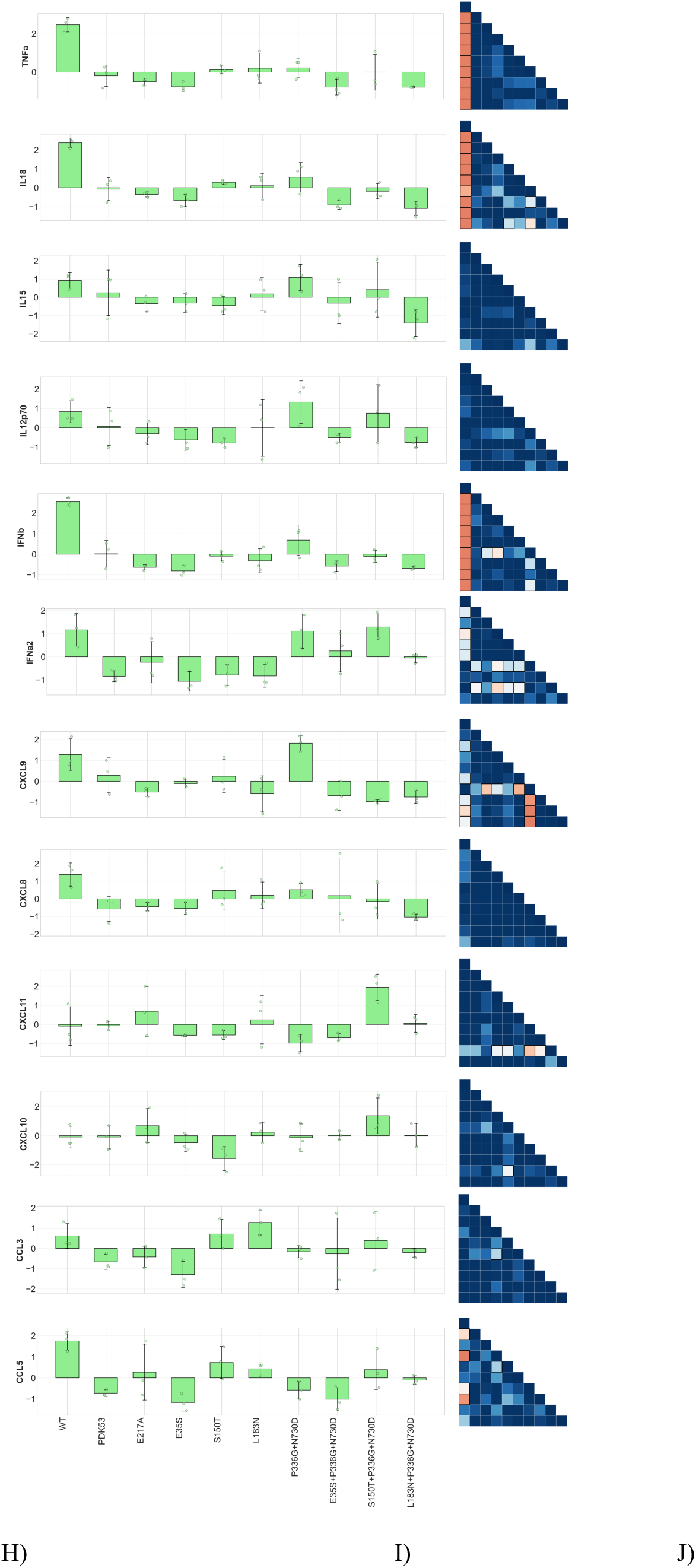
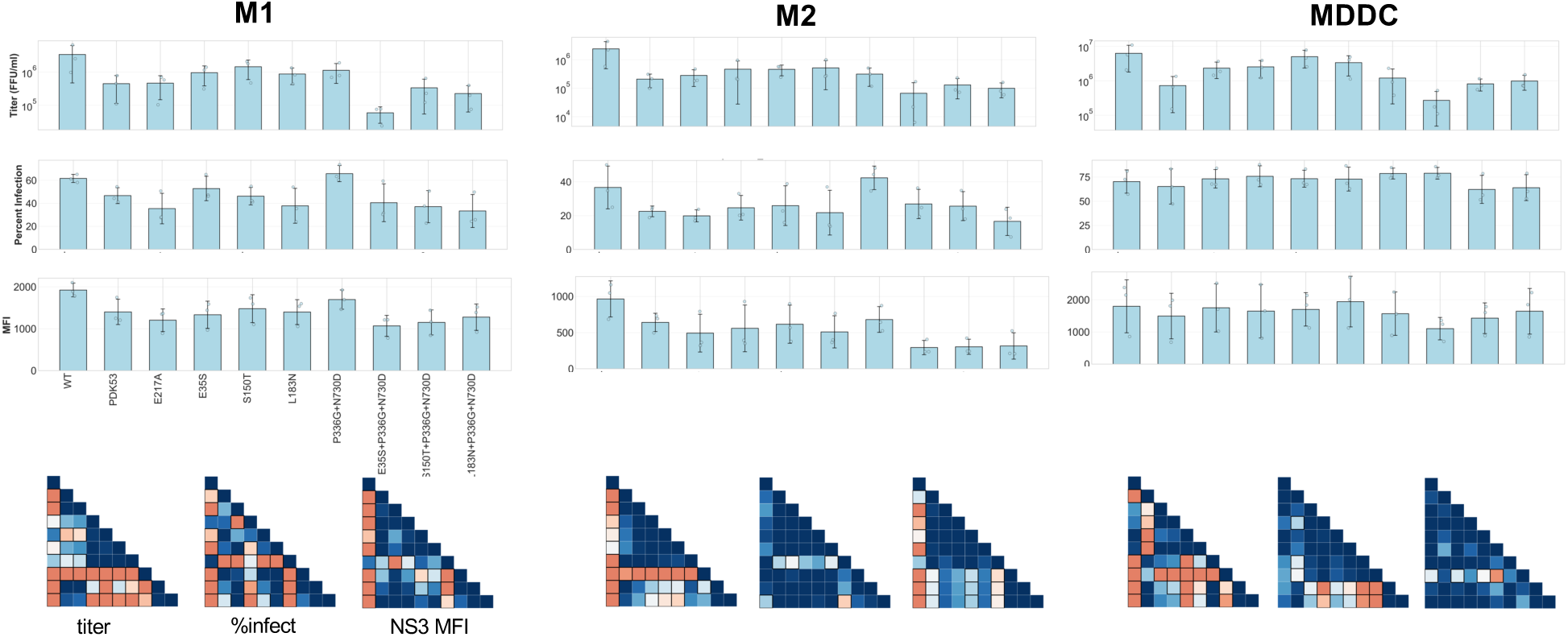
Comparison of immune-cell activation markers (A-D), cytokine/chemokine profiles (E-G), and infection phenotypes (H-J) across the DENV2 mutant panel in human MDDC (B,E,H), M1 (C,F,I), and M2 (D,G,J). A) The lower triangular heatmap visualizes the statistical significance of these pairwise comparisons, representing the -log_10_(adjusted *p*-value). Differences in marker levels among the viruses were assessed using a one-way ANOVA. Pairwise comparisons were performed using Tukey’s honestly significant difference (HSD) post-hoc test. Statistically significant pairs (*p* < 0.05) are highlighted with black square frames.

**Fig. S10:**
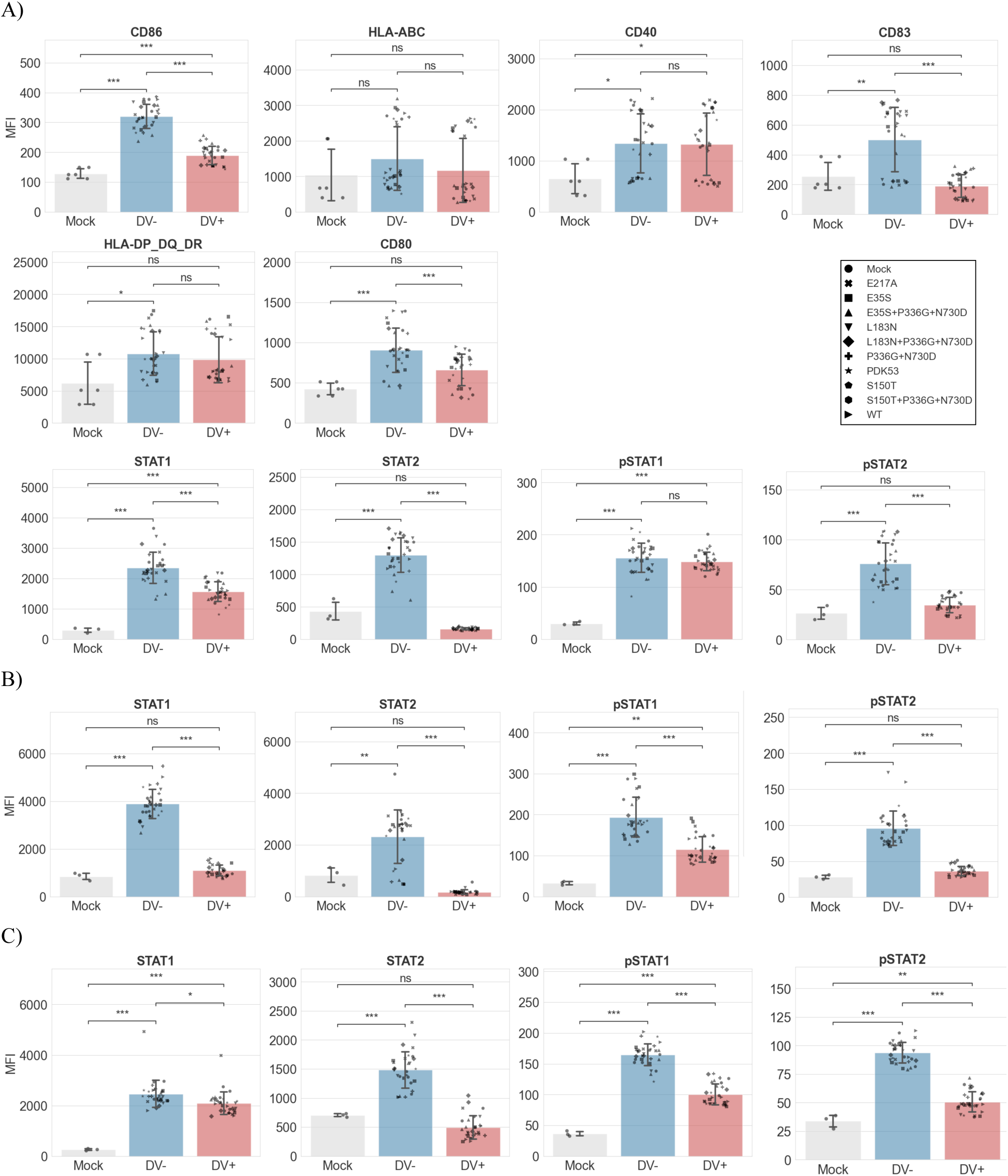
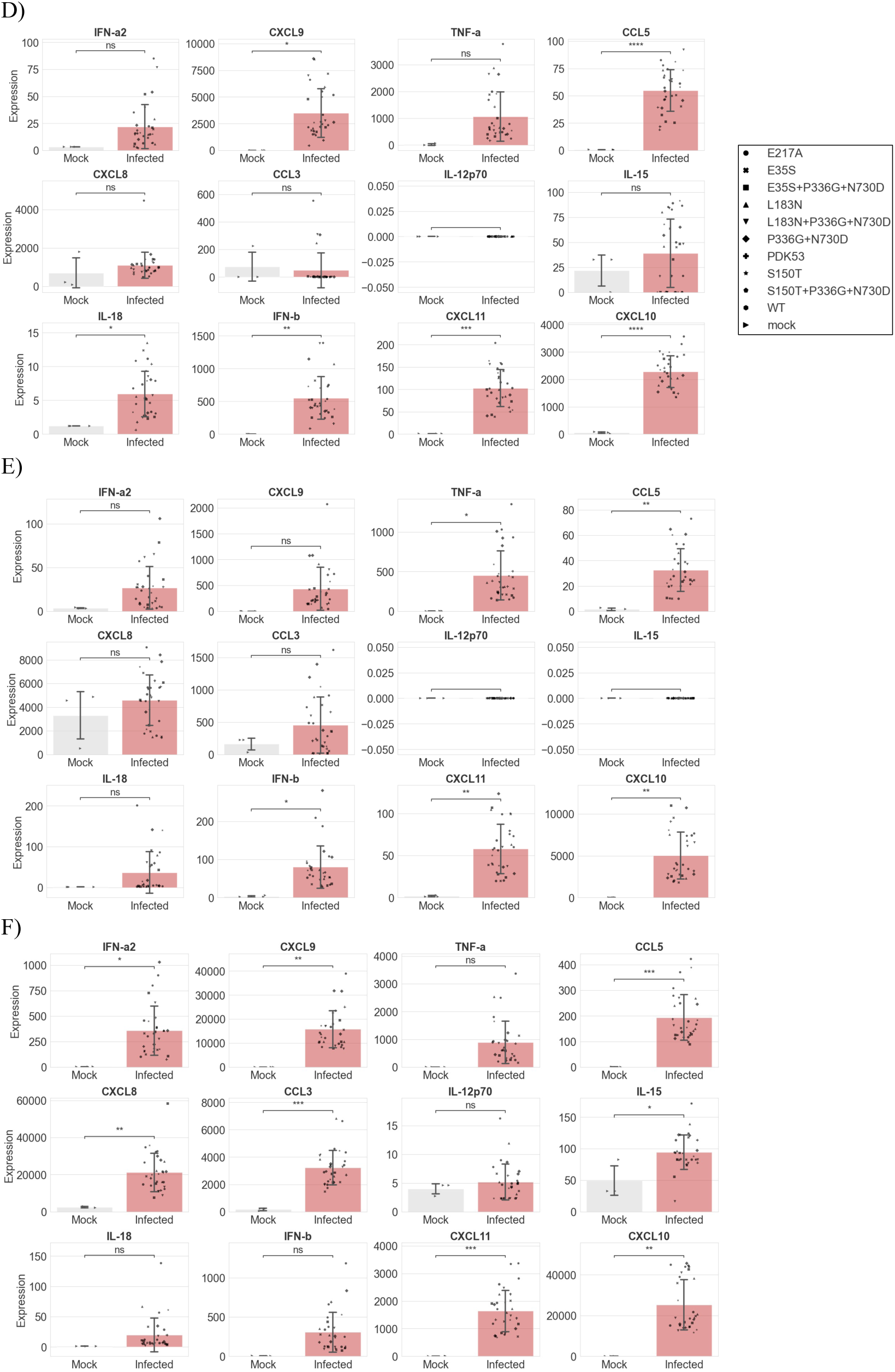
Comparison of immune-cell activation markers (A-C) and cytokine/chemokine profiles (D-F) of mock-infected, DENV-infected, and bystander human MDDC (A,D), M1 (B,E), and M2 (C,F). Statistical significance between groups was determined using an unpaired, two-tailed Student’s t-test. Significance is indicated as follows: ns (not significant), * *P* < 0.05, ** *P* < 0.01, *** *P* < 0.001, and **** *P* < 0.0001. Error bars represent the standard error of the mean (SEM).

## Notes

### Competing Interest Statement

Authors have filed patent applications relating to this work.

